# Interferon regulatory factor 6 drives hepatocyte redifferentiation during progenitor-driven liver regeneration in zebrafish

**DOI:** 10.64898/2026.11.23.768745

**Authors:** Nayoung Kim, Siyeo Lee, Tae-Young Choi

## Abstract

Severe liver injury induces biliary epithelial cells to dedifferentiate into proliferative hepatoblast-like liver progenitor cells (LPCs), which subsequently redifferentiate into mature hepatocytes; however, the intrinsic transcriptional regulators coordinating the exit from this progenitor state remain unclear. Here, we show that interferon regulatory factor 6 (*irf6*) is required for LPC redifferentiation during biliary-driven liver regeneration in zebrafish. By uncoupling maternal and zygotic functions in *irf6^wku2001^*mutants, we found that zygotic *irf6* is dispensable for primary liver organogenesis, whereas its loss significantly impairs hepatic mass recovery following hepatocyte ablation, coinciding with sustained hepatic *irf6* induction. Using a dual-fluorescent Notch reporter, *Tg(Tp1:VenusPEST);Tg(Tp1:H2BmCherry)*—where Notch attenuation (Venus^weak^/mCherry^+^) marks hepatocyte redifferentiation—we found that *irf6* is dispensable for generating Prox1^+^ LPCs. Instead, *irf6*-deficient transitioning cells expressing Hnf4α and betaine-homocysteine S-methyltransferase accumulate in a high-Notch (Venus^strong^/mCherry^+^) state and fail to efficiently transition into the Notch-attenuated (Venus^weak^/mCherry^+^) state. Crucially, *in vivo* signaling analyses demonstrated that *irf6* loss does not directly alter NF-κB or Wnt/beta-catenin activity within Notch-responsive LPCs, but selectively reduces NF-κB-active regenerating hepatocytes (CFP^+^/EGFP^+^) as a downstream consequence of impaired LPC-to-hepatocyte redifferentiation. Collectively, these findings indicate that *irf6* promotes hepatocyte redifferentiation by facilitating Notch signal attenuation during liver regeneration.

## Introduction

Interferon regulatory factor 6 (*irf6*) is a member of the interferon regulatory factor (Irf) family of transcription factors, and it exerts essential roles in epithelial development, cellular polarity, and tissue morphogenesis [1–3]. In humans, mutations in IRF6 cause congenital craniofacial disorders such as Van der Woude syndrome and popliteal pterygium syndrome, underscoring its critical function in epithelial differentiation and tissue integrity [2, 4]. At the cellular level, Irf6 regulates the balance between epithelial cell proliferation and terminal differentiation, facilitating cell cycle exit and the maintenance of epithelial architecture through transcriptional control of downstream targets [1, 5, 6].

During vertebrate development, *irf6* activity is tightly regulated both temporally and spatially. In zebrafish and Xenopus, *irf6* is required during early embryogenesis, particularly for primary superficial epithelial differentiation and gastrulation movements [7, 8]. Maternal-zygotic loss of *irf6* results in severe early developmental arrest and epithelial disintegration, which confirms its indispensable role during early embryogenesis [7, 8]; however, despite these well-established functions in embryonic epithelial tissues, the functional requirement for *irf6* at later developmental stages and in post-embryonic tissue contexts remains poorly understood. Notably, dynamic epithelial remodeling and cellular plasticity are defining features of post-injury organ regeneration [9]. Given its established role in orchestrating the epithelial proliferation-to-differentiation switch during embryogenesis, *irf6* may serve as an important transcriptional regulator of epithelial cell-state transitions in regenerative contexts; however, whether *irf6* participates in post-embryonic organ regeneration has not been systematically investigated.

In the liver of vertebrates, regeneration following tissue injury requires precise coordination of epithelial cell identity, proliferation, and differentiation [10]. Liver regeneration under mild-to-moderate injury conditions is largely driven by the compensatory self-duplication of pre-existing hepatocytes [11, 12], whereas severe hepatocyte loss can elicit an alternative regenerative program that relies on extensive biliary epithelial plasticity [13–15]. Under such extreme injury conditions, biliary epithelial cells (BECs) dedifferentiate into proliferative hepatoblast-like liver progenitor cells (LPCs), which subsequently redifferentiate into mature, functional hepatocytes [13–15]. Current studies on liver regeneration primarily focused on canonical extrinsic signaling pathways and broadly acting regulators, including Wnt/beta-catenin, Notch, Hippo–YAP, TGF-beta, and inflammatory NF-κB signaling [16–22]. A critical regulatory challenge during biliary-driven liver regeneration is the temporal resolution of progenitor-maintenance signals: although active Notch signaling is required to initiate BEC dedifferentiation and LPC expansion, Notch activity must subsequently be attenuated to permit LPCs to exit the progenitor state and redifferentiate into mature hepatocytes [14, 17, 23, 24]. Although extrinsic morphogen gradients guide these lineage transitions, the intrinsic epithelial transcription factors that facilitate Notch signal attenuation and drive terminal hepatocyte redifferentiation remain unclear.

Zebrafish constitute a suitable model system to address these questions, enabling genetic dissection of lineage transitions following targeted hepatocyte ablation. Using the nitroreductase/metronidazole (NTR/MTZ) system, mature hepatocytes can be selectively ablated to reliably induce a biliary-driven regenerative response [13, 25]. Furthermore, combining this ablation model with dual-fluorescent Notch reporter lines, *Tg(Tp1:VenusPEST);Tg(Tp1:H2BmCherry)*, allows real-time discrimination between Notch-active progenitors (Venus^strong^/mCherry^+^) and Notch-attenuated cells undergoing hepatocyte redifferentiation (Venus^weak^/mCherry^+^) at single-cell resolution.

In this study, we investigated the role of *irf6* during progenitor-driven liver regeneration in zebrafish. By generating a CRISPR/Cas9-induced loss-of-function mutant (*irf6^wku2001^*) to uncouple maternal and zygotic functions, we demonstrate that zygotic *irf6* is dispensable for primary liver organogenesis under homeostatic conditions, whereas it is specifically induced and required for efficient regenerative liver growth following severe hepatocyte ablation. Using *in vivo* lineage and signaling reporters, we found that *irf6* was not required for the initial generation of Prox1^+^ LPCs and did not directly alter progenitor-associated Wnt/beta-catenin or NF-κB signaling. Instead, loss of *irf6* caused transitioning cells expressing Hnf4α and betaine-homocysteine S-methyltransferase to accumulate in a high-Notch (Venus^strong^) state, impairing their transition into the Notch-attenuated (Venus^weak^) redifferentiation state and secondarily reducing the emergence of NF-κB-active regenerating hepatocytes. Collectively, our results identify *irf6* as a key intrinsic regulator that promotes hepatocyte redifferentiation by facilitating Notch signal attenuation during liver regeneration.

## Results

### Spatiotemporal expression patterns of *irf6* during zebrafish development

In our previous transcriptomic profiling across multiple stages of LPC-driven liver regeneration, we identified *irf6* as a candidate gene whose transcript levels were dynamically upregulated during the reparative process [26]. To establish the developmental baseline of *irf6* prior to evaluating its regenerative function, we characterized its spatiotemporal expression trajectory across embryonic and larval stages using RT-PCR and WISH analyses (Fig. 1). RT-PCR analysis demonstrated that *irf6* transcripts were abundantly detected from the two-cell stage through 50% epiboly, which reflected substantial maternal deposition, and remained continuously detectable throughout subsequent embryonic and larval development up to 5 dpf (Fig. 1A). Consistent with this temporal profile, WISH analysis revealed a broad, ubiquitous distribution of *irf6* transcripts across early cleavage and blastula stages (two-cell to 50% epiboly; Fig. 1B). From 24 hours post-fertilization (hpf) onward, coinciding with the onset of organogenesis, *irf6* expression became progressively compartmentalized to discrete anatomical regions, including the pharyngeal arches, tectum, hindbrain ventricle, pronephros, primitive gut, posterior somites, and optic vesicle (Fig. 1B). At later larval stages (3–5 dpf), *irf6* transcription was predominantly localized to endoderm-derived organs, including the liver, intestine, and swim bladder (Fig. 1B). Within these endodermal compartments, hepatic *irf6* expression was pronounced at 3 and 4 dpf before gradually decreasing by 5 dpf, establishing a spatial and temporal rationale for investigating *irf6* within the post-embryonic hepatic niche.

**Fig. 1.**
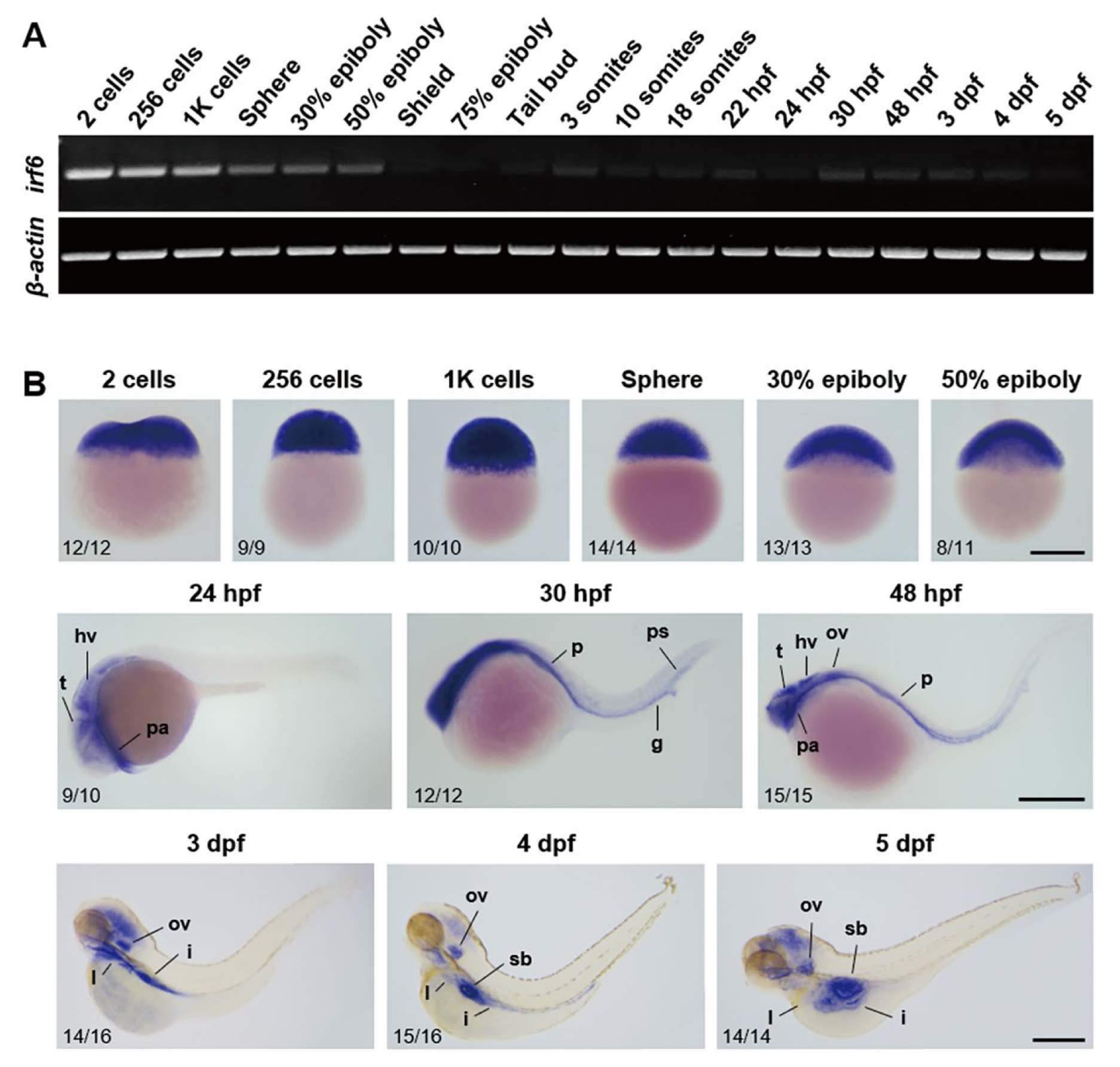
Spatiotemporal expression patterns of *irf6* during zebrafish development. (A) RT-PCR analysis of *irf6* transcript levels across developmental stages from the two-cell stage to 5 dpf. *irf6* transcripts were detected throughout embryonic and larval development. *β-actin* was used as an internal loading control. (B) WISH analysis of *irf6* expression patterns. During early cleavage and blastula stages (two-cell to 50% epiboly), *irf6* transcripts were broadly detected throughout the blastoderm. From 24 hours post-fertilization (hpf) onward, *irf6* expression became progressively restricted to specific anatomical regions, including the pharyngeal arches (pa), tectum (t), hindbrain ventricle (hv), pronephros (p), primitive gut (g), posterior somites (ps), and optic vesicle (ov). At larval stages (3–5 dpf), *irf6* expression was predominantly observed in endoderm-derived organs, including the liver (l), intestine (i), and swim bladder (sb). Numbers in the bottom-left corner of each panel indicate the number of embryos or larvae exhibiting the representative expression pattern out of the total number examined. Scale bars: 250 μm.

### Generation of the *irf6^wku2001^* loss-of-function allele and developmental uncoupling of maternal and zygotic *irf6* requirements

To investigate the endogenous requirement for *irf6*, we generated the *irf6^wku2001^* mutant allele using CRISPR/Cas9-mediated genome editing targeted to the coding region of the *irf6* locus (Fig. 2A). Sequence analysis of the recovered allele identified a 23-bp deletion within the coding sequence, which induced a frameshift and introduced a premature stop codon (Fig. 2B). *In silico* translation predicted that the *irf6^wku2001^* allele would produce a truncated 22-amino acid peptide, compared with the full-length 492-amino acid wild-type Irf6 protein (Fig. 2C). This truncation eliminated all major functional regions, including the winged-helix DNA-binding domain and the SMAD/FHA domain (Fig. 2A, C). PCR-based genotyping using gene-specific primers further confirmed the 23-bp deletion, yielding a 157-bp amplicon in *irf6^wku2001^* mutants compared with the 180-bp wild-type band (Fig. 2D). Collectively, these structural alterations indicate that *irf6^wku2001^* operates as a loss-of-function allele predicted to abolish Irf6 protein function.

**Fig. 2.**
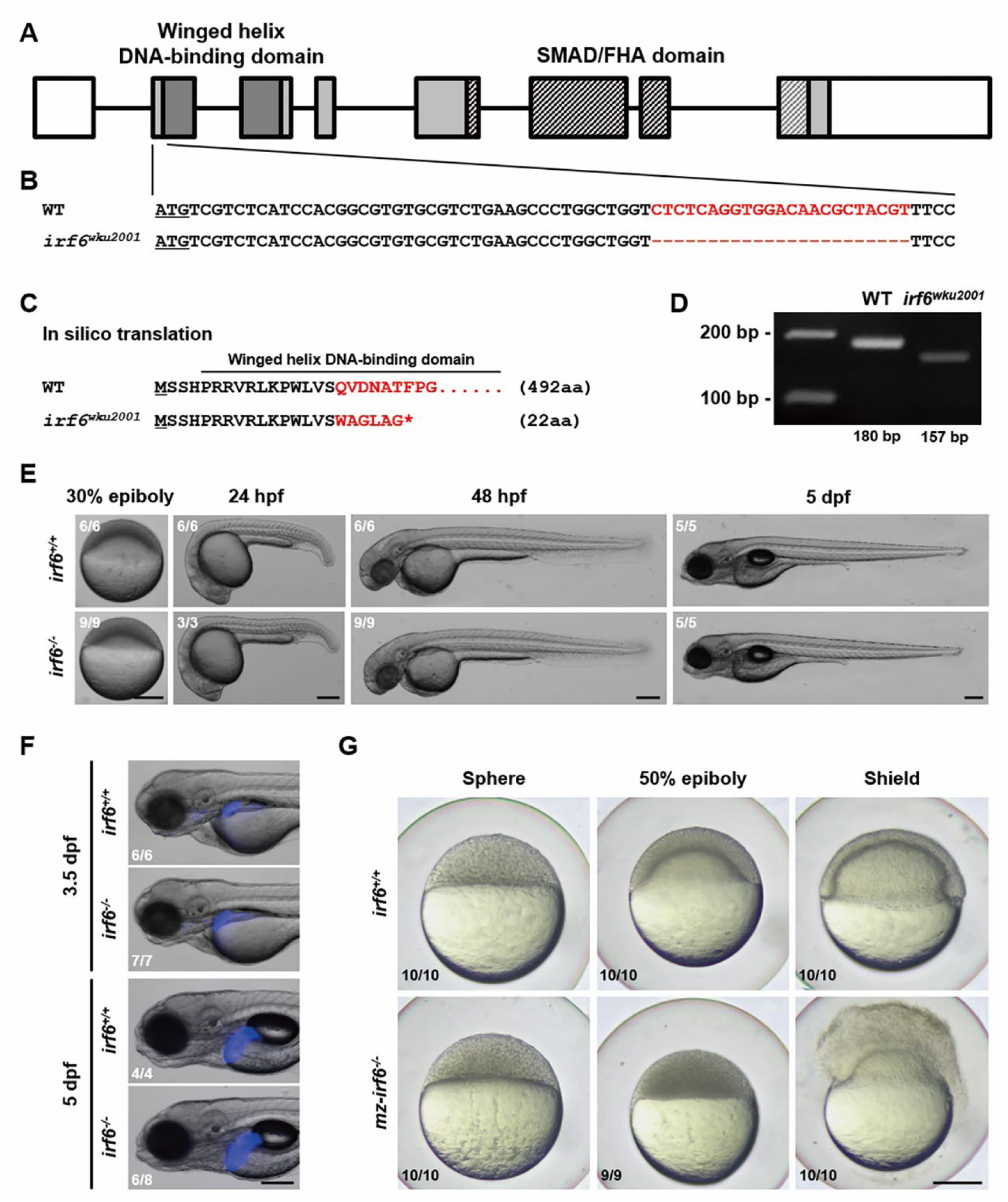
Generation of the *irf6^wku2001^* mutant allele and morphological analysis of developing embryos. (A) Schematic representation of the *irf6* genomic locus and corresponding Irf6 protein functional domains. White boxes indicate untranslated regions (UTRs), light gray boxes represent coding exon regions, dark gray boxes denote the winged-helix DNA-binding domain, and striped boxes indicate the SMAD/FHA domain. (B) Nucleotide sequence alignment of wild-type (WT) and *irf6^wku2001^*mutant alleles, showing a 23-bp deletion (indicated in red in WT) within the coding region that induces a frameshift mutation. The translation initiation codon (ATG) is underlined. (C) *In silico* translation demonstrating that the 23-bp deletion introduces a premature stop codon (asterisk), producing a truncated peptide of 22 amino acids (aa) lacking all functional domains, compared to the full-length WT protein (492 aa). Altered amino acid residues are shown in red. (D) PCR-based genotyping of WT and *irf6^wku2001^* mutant embryos. The WT allele yielded a 180-bp amplicon, whereas the mutant allele yielded a 157-bp amplicon, confirming the 23-bp deletion. (E) Bright-field microscopy images comparing *irf6^+/+^* and zygotic *irf6^-/-^* embryos at 30% epiboly and lateral views at 24 hpf, 48 hpf, and 5 dpf, showing comparable overall morphology. (F) Lateral fluorescence images merged with bright-field views of *Tg(fabp10a:*CFP-NTR) larvae at 3.5 dpf and 5 dpf, showing comparable CFP-positive hepatocyte signals in the liver (blue) between *irf6^+/+^* and *irf6^-/-^* larvae. (G) Bright-field images comparing *irf6^+/+^* and maternal-zygotic *irf6^wku2001^* (mz-*irf6^-/-^*) mutant embryos at early developmental stages (sphere, 50% epiboly, and shield stages). While *irf6^+/+^* embryos progressed normally through epiboly, mz-*irf6^-/-^* mutants failed to develop beyond the sphere stage and exhibited severe morphological disintegration at the shield stage. Numbers in the corners of panels (E–G) indicate the number of embryos or larvae exhibiting the representative phenotype out of the total number examined. Scale bars: 250 μm.

To determine whether zygotic *irf6* is required for normal embryonic development and primary hepatogenesis, we evaluated the morphological progression of *irf6^+/+^*, *irf6^+/-^*, and zygotic *irf6^-/-^* mutant embryos. Zygotic *irf6^-/-^* mutants progressed through primary developmental milestones without overt morphological abnormalities from 30% epiboly through 24 hpf, 48 hpf, and 5 dpf (Fig. 2E). Structural evaluation using the hepatocyte-specific reporter line *Tg(fabp10a:*CFP-NTR) demonstrated that primary liver specification and parenchymal expansion were morphologically comparable across genotypes, with equivalent CFP-positive hepatocyte signals observed in *irf6^+/+^* and *irf6^-/-^* larvae at both 3.5 and 5 dpf (Fig. 2F). In contrast, maternal-zygotic *irf6^wku2001^* (mz-*irf6^-/-^*) mutant embryos exhibited severe early developmental arrest (Fig. 2G). Wild-type control embryos progressed normally through blastula and gastrula stages, whereas mz-*irf6^-/-^* mutants failed to develop beyond the sphere stage and displayed pronounced morphological disintegration at the shield stage (Fig. 2G). These dynamic developmental defects were further confirmed by time-lapse imaging (Supplementary Video 1), which closely recapitulates the early epiboly arrest and superficial epithelial disintegration phenotypes previously reported in maternal and maternal-zygotic *irf6*-deficient zebrafish models [7, 8]. This phenotypic concordance functionally validates the loss-of-function nature of our newly generated *irf6^wku2001^* allele *in vivo* and confirms an essential requirement for maternally supplied *irf6* during early embryogenesis. Taken together, this developmental uncoupling demonstrates that while maternal *irf6* is strictly required for early embryonic progression, zygotic *irf6* is dispensable for initial liver organogenesis, thereby providing a structurally intact hepatic baseline for subsequent post-embryonic regeneration analyses.

### Loss of zygotic *irf6* impairs expansion of the regenerating liver following targeted hepatocyte ablation

Given the expression of *irf6* in the larval liver and the structurally intact hepatic baseline observed in zygotic *irf6^-/-^* mutants, we next investigated whether *irf6* is required for LPC-driven liver regeneration following severe parenchymal injury. Targeted hepatocyte ablation was induced using the NTR/Mtz system in *Tg(fabp10a:CFP-NTR);Tg(fabp10a:*DsRed) double-transgenic larvae from 3.5 to 5 dpf (ablation 0 h [A0h] to A36h), followed by Mtz washout and morphological and quantitative analyses at defined regeneration time points: 24, 48, and 72 hours post-washout (A36h-R24h, A36h-R48h, and A36h-R72h; Fig. 3A, Supplementary Fig. 1A). Following hepatocyte ablation, regenerating DsRed-positive liver tissue was observed across *irf6^+/+^*, *irf6^+/-^*, and *irf6^-/-^* mutant larvae throughout the recovery phase (Fig. 3B, Supplementary Fig. 1B). Quantitative morphometric analysis demonstrated that heterozygous (*irf6^+/-^*) larvae exhibited regenerating liver areas comparable to those of wild-type (*irf6^+/+^*) controls across all examined time points, confirming that a single functional allele of *irf6* is sufficient to maintain normal regenerative capacity (Supplementary Fig. 1B, C). In contrast, homozygous *irf6^-/-^* mutant larvae displayed a significantly reduced regenerating liver area compared with *irf6^+/+^* controls at A36h-R24h, and this reduction persisted through A36h-R48h and A36h-R72h (Fig. 3B, C). Collectively, these results indicate that loss of zygotic *irf6* specifically impairs compensatory regenerative liver growth rather than initial liver formation.

**Fig. 3.**
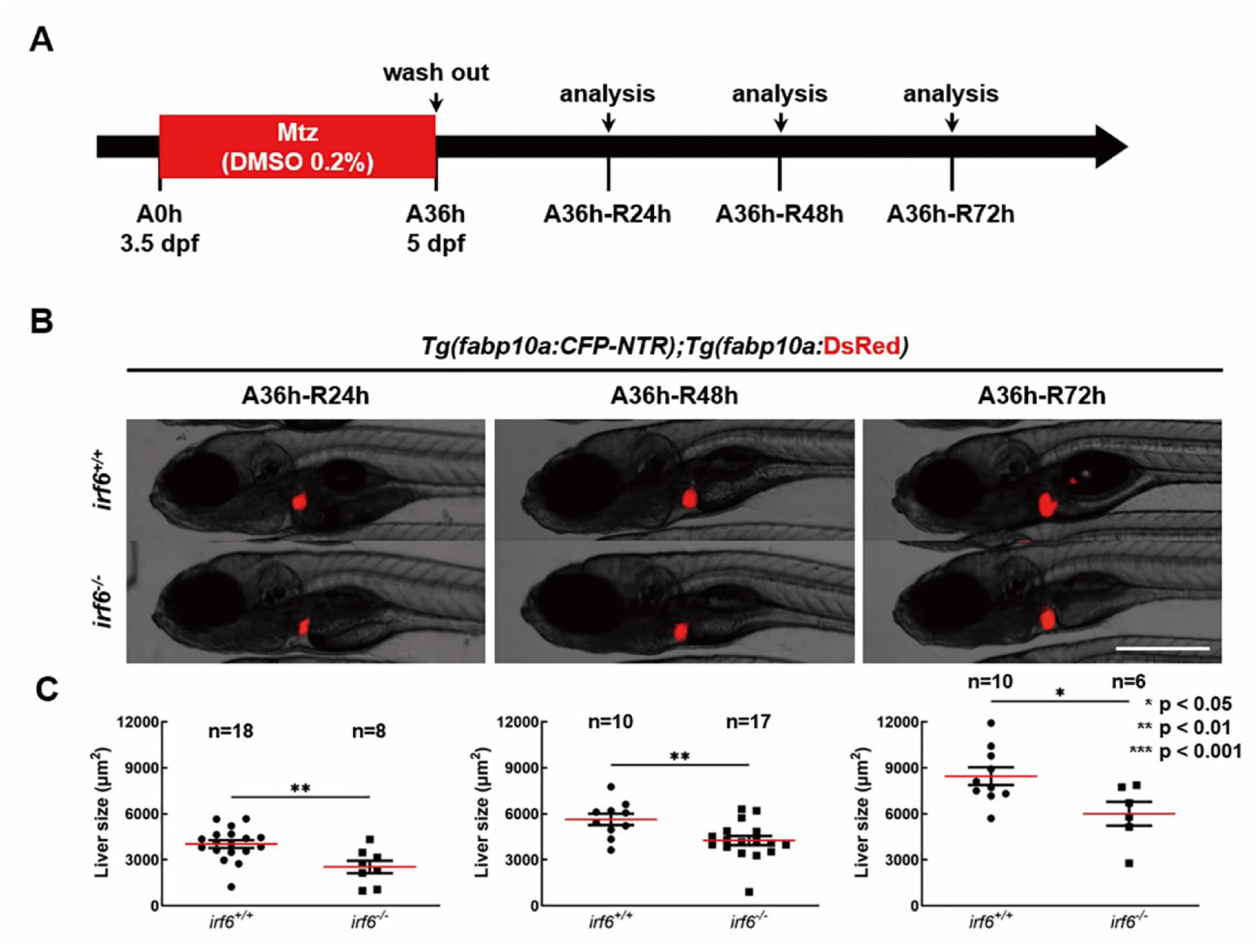
Reduced liver size in the *irf6^wku2001^* mutant during liver regeneration. (A) Experimental scheme illustrating the timeline of metronidazole (Mtz; in 0.2% DMSO) treatment for targeted hepatocyte ablation from 3.5 to 5 dpf (A0h to A36h), followed by Mtz washout and subsequent analysis at defined regeneration time points (A36h-R24h, A36h-R48h, and A36h-R72h). (B) Representative lateral fluorescence images merged with bright-field views showing regenerating livers (DsRed, red) at A36h-R24h, A36h-R48h, and A36h-R72h in *irf6^+/+^* and *irf6^- /-^*larvae carrying the *Tg(fabp10a:CFP-NTR);Tg(fabp10a:*DsRed) transgenes. (C) Quantification of regenerating liver size (μm^2^) at A36h-R24h (left), A36h-R48h (middle), and A36h-R72h (right), measured from DsRed-positive fluorescence images. Each data point represents an individual larva, and the number of examined larvae (n) is indicated above each group. Red horizontal bars indicate the mean, and black error bars indicate the standard deviation (SD). Statistical significance was determined using an unpaired two-tailed Student’s t test (\**P < 0.05*; \*\**P < 0.01*; \*\*\**P < 0.001*; ns., not significant). Scale bar: 500 μm.

### *irf6* expression is dynamically sustained during liver regeneration alongside preserved whole-mount expression of sequential lineage regulators

To determine whether *irf6* transcription was actively recruited during injury-induced hepatic repair, we performed WISH analysis across defined ablation and regeneration stages (Fig. 4A). Basal *irf6* expression in the liver had largely subsided in unablated control larvae between 5 and 7 dpf; however, *irf6* transcripts were distinctly induced and maintained within the regenerating hepatic region during late ablation (A33h and A36h) and persisted throughout the early and intermediate phases of regeneration (A36h-R6h, A36h-R24h, and A36h-R48h; Fig. 4B, red arrows). This sustained post-injury expression, which corroborated our initial transcriptomic profiling, indicated that *irf6* was actively reactivated within the hepatic niche during the reparative response.

**Fig. 4.**
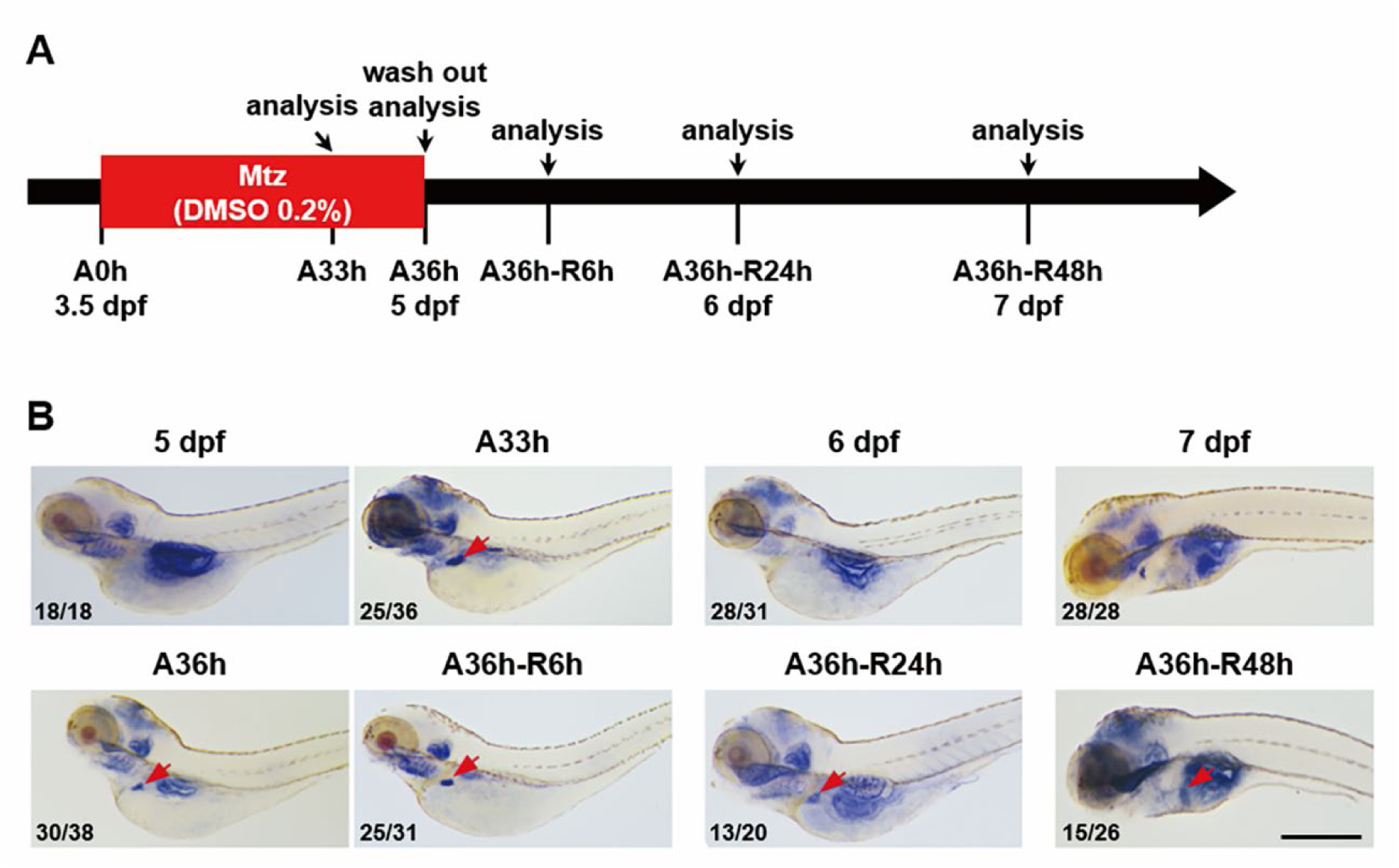
Dynamic *irf6* expression during liver regeneration. (A) Experimental scheme illustrating Mtz-induced (in 0.2% DMSO) hepatocyte ablation from 3.5 to 5 dpf (A0h to A36h), followed by Mtz washout and WISH analysis at the indicated time points during ablation and regeneration (A33h, A36h, A36h-R6h, A36h-R24h [6 dpf], and A36h-R48h [7 dpf]). (B) Representative lateral WISH images showing *irf6* expression patterns in unablated control larvae (5, 6, and 7 dpf) and Mtz-treated larvae at the indicated stages. *irf6* expression was distinctly detected in the regenerating liver region at A33h and A36h and persisted throughout liver regeneration at A36h-R6h, A36h-R24h, and A36h-R48h (red arrows), whereas hepatic *irf6* transcripts were barely detectable in age-matched unablated controls. Numbers in the bottom-left corner of each panel indicate the number of larvae exhibiting the representative expression pattern out of the total number examined. Scale bar: 250 μm.

To evaluate whether the regenerative growth defect in *irf6^-/-^* mutants stemmed from a global failure in hepatic fate acquisition at the whole-tissue level, we next examined the spatiotemporal expression of key developmental transcription factors and mature hepatocyte functional markers across *irf6^+/+^*, *irf6^+/-^*, and *irf6^-/-^* larvae using WISH (Supplementary Fig. 2 and Supplementary Fig. 3). We first profiled four core transcription factors governing sequential steps of hepatic lineage progression across late ablation and early regeneration stages (A33h, A36h, A36h-R6h, and A36h-R24h; Supplementary Fig. 2A, B). Specifically, we examined *foxa3* and *hhex*, which regulate sequential stages along the foregut endoderm-to-liver trajectory spanning hepatic competence, specification, and liver bud morphogenesis, as well as *prox1* and *hnf4α*, which act as cooperative yet contrasting transcriptional regulators that balance hepatoblast/progenitor maintenance and hepatocyte differentiation. Across late ablation and early regeneration stages (A33h to A36h-R24h), both the competence/specification factors (*foxa3* and *hhex*) and the progenitor maintenance/differentiation regulators (*prox1* and *hnf4α*) were comparably induced in the regenerating liver bud of *irf6^+/+^*, *irf6^+/-^*, and *irf6^-/-^* larvae (Supplementary Fig. 2A, B). Furthermore, mature hepatocyte markers—including *lfabp*, *gc*, *cp*, and *bhmt*—were expressed in the regenerating liver across all genotypes from A36h-R24h to A36h-R48h, although the stained hepatic domain appeared smaller in *irf6^-/-^* mutants, consistent with their reduced regenerating liver size (Supplementary Fig. 3A, B). Collectively, the preserved whole-mount expression of these sequential transcriptional regulators and mature hepatocyte markers indicated that *irf6* deficiency did not abrogate initial biliary activation or global hepatic specification, which prompted us to examine how the balance between progenitor maintenance (Prox1) and hepatocyte differentiation (Hnf4α and Bhmt) was resolved at single-cell resolution.

### *irf6* loss alters the distribution of Notch-responsive transitional cell states during LPC-driven hepatocyte redifferentiation

During LPC-driven liver regeneration following severe hepatocyte ablation, BECs dedifferentiate into proliferative hepatoblast-like LPCs expressing both Prox1 and Hnf4α, which subsequently redifferentiate into mature hepatocytes [14, 26]. This lineage progression is tightly governed by dynamic changes in Notch signaling: high Notch activity is required to maintain BECs and early transitional LPCs, whereas the subsequent attenuation of Notch signaling is strictly required for LPCs to exit the progenitor state and redifferentiate into hepatocytes. To assess how *irf6* regulated cell fate transitions along this Notch-dependent trajectory, we performed immunofluorescence analyses using the dual-fluorescent Notch reporter line *Tg(Tp1:*VenusPEST*);Tg(Tp1:*H2BmCherry) at late ablation (A33h) and early regeneration (A36h-R6h) stages (Fig. 5A and Supplementary Fig. 4A). Because H2BmCherry is highly stable while VenusPEST is rapidly degraded, this system distinguishes cells maintaining active Notch signaling (Venus^strong^ and mCherry^+^) from biliary-derived cells that have downregulated Notch signaling (Venus^weak^ and mCherry^+^). During LPC-driven liver regeneration, Venus^strong^ and mCherry^+^ cells represent BECs or uncommitted LPCs retaining active Notch signaling, whereas Venus^weak^ and mCherry^+^ cells represent transitional cells actively undergoing redifferentiation into hepatocytes following Notch signal attenuation. Therefore, we first examined the expression of Prox1, a core transcription factor required for LPC establishment and hepatic lineage maintenance, within the Tp1-positive population (Fig. 5B and Supplementary Fig. 4B). At A33h, although the total numbers of Venus^+^/mCherry^+^ and Prox1^+^/Venus^+^/mCherry^+^ cells were transiently lower in *irf6^-/-^* mutants than in *irf6^+/-^*controls (Supplementary Fig. 4B), virtually all Venus^+^/mCherry^+^ cells expressed Prox1 in both genotypes (∼100%; Fig. 5B). By A36h-R6h, the total numbers of Venus^+^/mCherry^+^ and Prox1^+^/Venus^+^/mCherry^+^ cells recovered to levels comparable to controls (Supplementary Fig. 4B), whereas the percentage of Prox1^+^/Venus^+^/mCherry^+^ cells among total Venus^+^/mCherry^+^ cells was modestly but significantly decreased in *irf6^-/-^* mutants (Fig. 5B). Stratification of Tp1-positive cells by Venus fluorescence intensity and Prox1 expression (Prox1^+^ vs. Prox1^-^) revealed that while both the number and relative proportion of the redifferentiating Venus^weak^/mCherry^+^ population remained comparable between genotypes, the absolute number of Prox1^+^/Venus^strong^/mCherry^+^ cells was significantly reduced in *irf6^-/-^* mutants at A33h (Fig. 5B and Supplementary Fig. 4B). These results indicated that Prox1-positive LPCs were robustly generated in *irf6^-/-^* mutants, although their abundance and maintenance within the high-Notch compartment were subtly altered during the transition from late ablation to early regeneration.

**Fig. 5.**
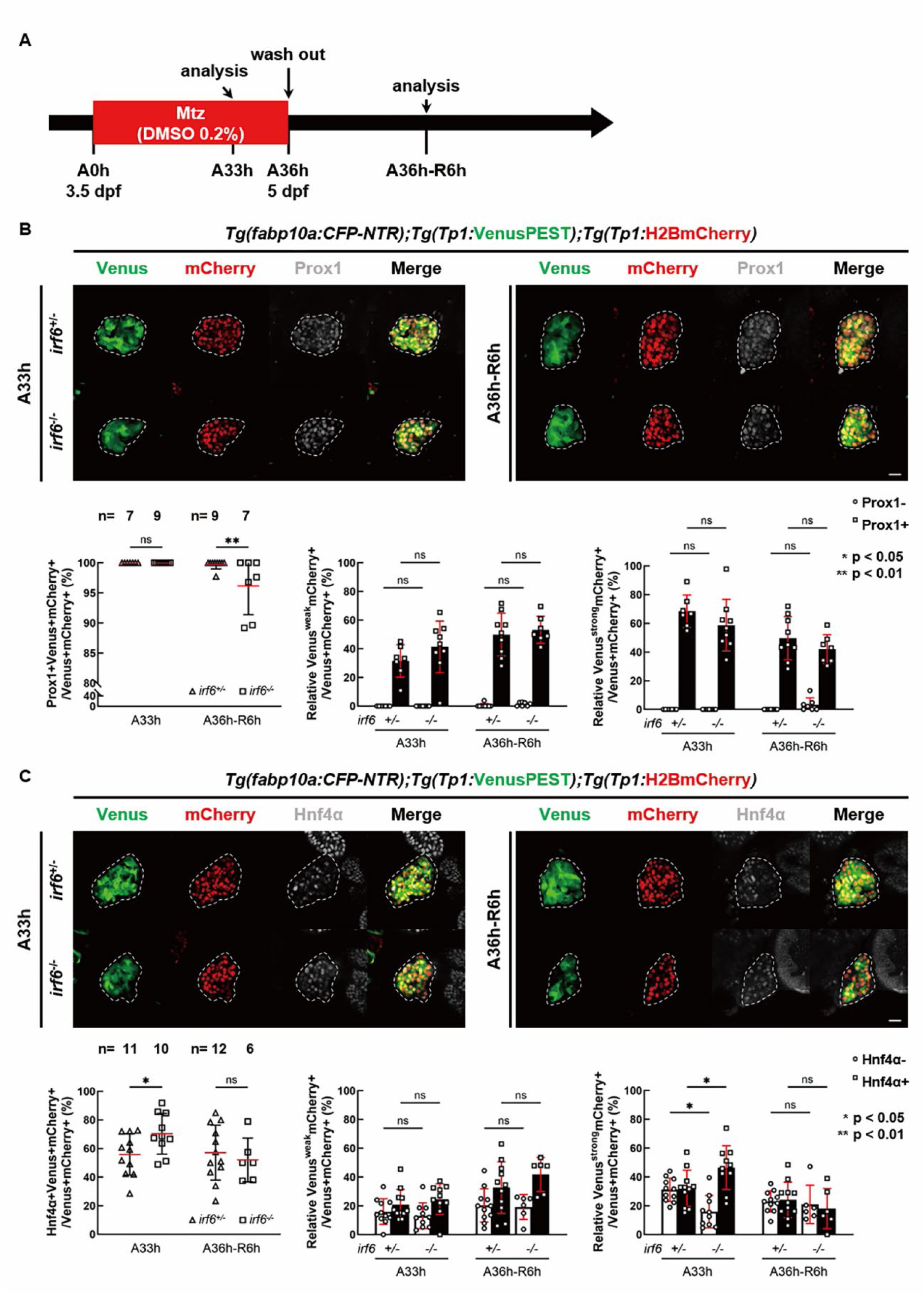
Quantitative immunofluorescence analysis of Prox1 and Hnf4*α* expression across Notch-responsive cell states during liver regeneration in *irf6* mutant larvae. (A) Experimental scheme illustrating Mtz-induced (in 0.2% DMSO) hepatocyte ablation from 3.5 to 5 dpf (A0h to A36h) and the time points analyzed during late ablation (A33h) and early liver regeneration (A36h-R6h). (B) Representative confocal immunofluorescence images (top) and quantitative analyses (bottom) of regenerating livers from *irf6^+/-^* control and *irf6^-/-^* mutant larvae at A33h and A36h-R6h using the *Tg(fabp10a:CFP-NTR);Tg(Tp1:*VenusPEST*);Tg(Tp1:*H2BmCherry) reporter line immunostained for Prox1 (gray, marking hepatic progenitor-associated nuclear identity). Unstable VenusPEST (green) and stable H2BmCherry (red) label Tp1-positive Notch-responsive biliary-derived cells, with Venus intensity distinguishing Venus^strong^ and Venus^weak^ subpopulations. Graphs show the percentage of Prox1^+^/Venus^+^/mCherry^+^ cells among total Venus^+^/mCherry^+^ cells (left) and the relative percentages of Venus^weak^/mCherry^+^ (middle) and Venus^strong^/mCherry^+^ (right) cells stratified by Prox1 expression (Prox1^-^, open circles/white bars; Prox1^+^, open squares/black bars). (C) Representative confocal immunofluorescence images (top) and quantitative analyses (bottom) of regenerating livers from *irf6^+/-^* and *irf6^-/-^* larvae at A33h and A36h-R6h immunostained for Hnf4α (gray, marking hepatoblast/hepatocyte nuclear identity). Graphs show the percentage of Hnf4α^+^/Venus^+^/mCherry^+^ cells among total Venus^+^/mCherry^+^ cells (left) and the relative percentages of Venus^weak^/mCherry^+^ (middle) and Venus^strong^/mCherry^+^ (right) cells stratified by Hnf4α expression (Hnf4α^-^, open circles/white bars; Hnf4α^+^, open squares/black bars). Dashed lines outline the regenerating liver. Each data point represents an individual larva, with the number of examined larvae (n) indicated above the graphs. Red bars indicate the mean ± SD. Statistical significance was determined using an unpaired two-tailed Student’s t test (\**P < 0.05*; \*\**P < 0.01*; \*\*\**P < 0.001*; ns., not significant). Scale bars: 20 μm.

We next analyzed the expression of Hnf4α, a definitive transcriptional driver of hepatoblast specification and hepatocyte identity acquisition, to evaluate whether LPCs properly transition from the Notch-active (Venus^strong^) state to the Notch-attenuated (Venus^weak^) redifferentiation state (Fig. 5C and Supplementary Fig. 4C). While the total numbers of Venus^+^/mCherry^+^ cells and Hnf4α^+^/Venus^+^/mCherry^+^ cells were comparable between *irf6^+/-^* control and *irf6^-/-^* mutant larvae at both A33h and A36h-R6h (Supplementary Fig. 4C), the overall percentage of Hnf4α^+^/Venus^+^/mCherry^+^ cells among Venus^+^/mCherry^+^ cells was modestly elevated in *irf6^-/-^*mutants at A33h (Fig. 5C). Subdivision of the Tp1-positive pool into Venus^strong^ and Venus^weak^ states combined with Hnf4α expression (Hnf4α^+^ vs. Hnf4α^-^) showed a pronounced shift in cell-state distribution at A33h (Fig. 5C and Supplementary Fig. 4C). In *irf6^-/-^* mutants, both the absolute number and relative proportion of Hnf4α^+^/Venus^strong^/mCherry^+^ cells were significantly increased at A33h, accompanied by a reciprocal decrease in the Hnf4α^-^/Venus^strong^/mCherry^+^ population (Fig. 5C and Supplementary Fig. 4C). Despite this accumulation of Hnf4α-expressing cells within the Venus^strong^ compartment, the number and relative proportion of Hnf4α^+^/Venus^weak^/mCherry^+^ cells—representing cells that have successfully attenuated Notch signaling to undergo hepatocyte redifferentiation—failed to increase correspondingly (Fig. 5C and Supplementary Fig. 4C). Collectively, these findings indicate that while *irf6* is dispensable for initial LPC induction, loss of *irf6* causes Hnf4α-positive transitional cells to transiently accumulate in a high-Notch Venus^strong^ state, suggesting an impaired or delayed transition toward the Notch-attenuated Venus^weak^ state required for hepatocyte redifferentiation.

### *irf6* is required for Notch signal attenuation during the redifferentiation of Tp1-positive progenitors into Bhmt-positive mature hepatocytes

To determine whether the impaired transition from the high-Notch (Venus^strong^) state to the Notch-attenuated (Venus^weak^) state directly compromised terminal hepatocyte redifferentiation, we examined the expression of Bhmt, a definitive marker of mature, metabolically functional hepatocytes, at A33h and A36h-R6h (Fig. 6A, B, Supplementary Fig. 5A, B). Quantitative analysis revealed that the total number of Bhmt^+^/Venus^+^/mCherry^+^ cells was comparable between *irf6^+/-^* control and *irf6^-/-^*mutant larvae at both A33h and A36h-R6h (Supplementary Fig. 5B). In contrast, the total number of Venus^+^/mCherry^+^ cells was transiently reduced in *irf6* mutants at A33h—resulting in a higher apparent percentage of Bhmt^+^/Venus^+^/mCherry^+^ cells among the total Venus^+^/mCherry^+^ pool at this time point—before recovering to control levels by A36h-R6h (Fig. 6B, Supplementary Fig. 5B).

**Fig. 6.**
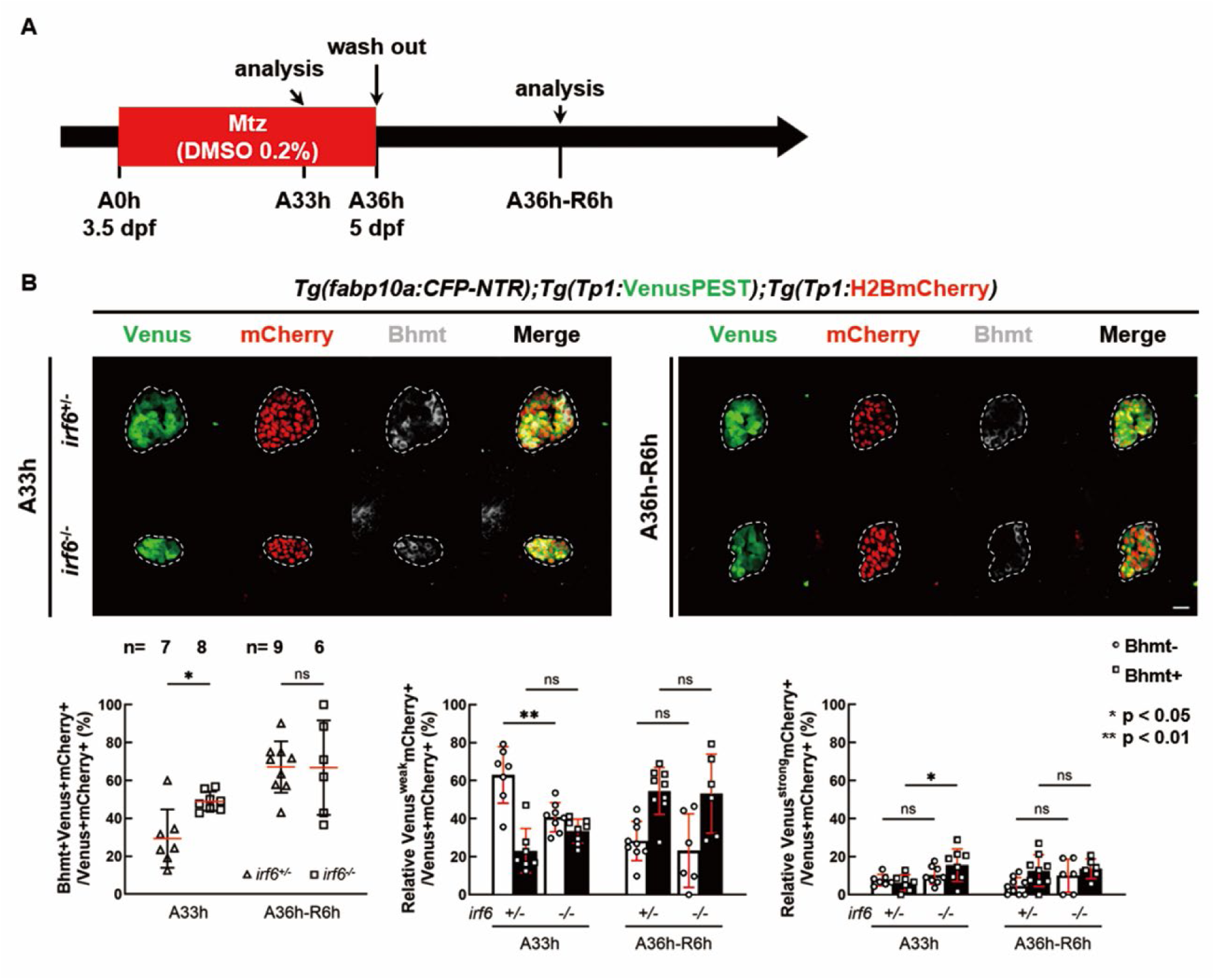
Quantitative immunofluorescence analysis of mature hepatocyte marker (Bhmt) expression during liver regeneration in *irf6* mutant larvae. (A) Experimental scheme illustrating Mtz-induced (in 0.2% DMSO) hepatocyte ablation from 3.5 to 5 dpf (A0h to A36h) and the time points analyzed during late ablation (A33h) and early liver regeneration (A36h-R6h). (B) Representative confocal immunofluorescence images (top) and quantitative analyses (bottom) of regenerating livers from *irf6^+/-^* control and *irf6^-/-^* mutant larvae at A33h and A36h-R6h using the *Tg(fabp10a:CFP-NTR);Tg(Tp1:*VenusPEST*);Tg(Tp1:*H2BmCherry) reporter line immunostained for Bhmt (gray, marking differentiated hepatocytes). Unstable VenusPEST (green) and stable H2BmCherry (red) label Tp1-positive Notch-responsive biliary-derived cells, with Venus fluorescence intensity distinguishing high-Notch (Venus^strong^) progenitors from Notch-attenuated (Venus^weak^) redifferentiating cells. Graphs show the percentage of Bhmt^+^/Venus^+^/mCherry^+^ cells among total Venus^+^/mCherry^+^ cells (left) and the relative percentages of Venus^weak^/mCherry^+^ (middle) and Venus^strong^/mCherry^+^ (right) populations among total Venus^+^/mCherry^+^ cells subdivided into Bhmt^-^(white bars, circles) and Bhmt^+^ (black bars, squares) fractions. Dashed lines outline the regenerating liver. Each data point represents an individual larva, with the number of examined larvae (n) indicated above the graphs. Red bars indicate the mean **±** SD. Statistical significance was determined using an unpaired two-tailed Student’s t test (\**P < 0.05*; \*\**P < 0.01*; \*\*\**P < 0.001*; ns., not significant). Scale bar: 20 μm.

To evaluate how efficiently Tp1-derived cells undergo terminal hepatocyte redifferentiation, we subdivided the Tp1-positive population into Notch-attenuated (Venus^weak^/mCherry^+^) and Notch-active (Venus^strong^/mCherry^+^) states in conjunction with Bhmt expression (Fig. 6B, Supplementary Fig. 5B). In control larvae undergoing LPC-driven liver regeneration, as transitioning cells extinguished Notch signaling to redifferentiate into mature hepatocytes, Bhmt expression became progressively enriched within the Venus^weak^/mCherry^+^ population from A33h to A36h-R6h (Fig. 6B, Supplementary Fig. 5B). In *irf6* mutants, however, this differentiation-coupled shift toward the Venus^weak^ state was disrupted at A33h: both the absolute number and relative proportion of Bhmt^-^/Venus^weak^/mCherry^+^ cells entering the Notch-attenuated state were significantly reduced, whereas the relative proportion of Bhmt^+^/Venus^strong^/mCherry^+^ cells was significantly elevated compared with *irf6^+/-^* controls (Fig. 6B, Supplementary Fig. 5B). This enrichment of Bhmt^+^/Venus^strong^/mCherry^+^ cells indicated that *irf6*-deficient transitional cells initiated mature marker expression while retaining active Notch signaling, skewing the Tp1-derived population toward a persistent Venus^strong^ state. Taken together with our analyses of Prox1 and Hnf4α (Fig. 5), these findings demonstrate that *irf6* is dispensable for the initial generation of Tp1-positive Notch-responsive LPCs following severe hepatocyte ablation but is required for their efficient progression through redifferentiation-associated cell states. Loss of *irf6* disrupts the timely transition from the Notch-active (Venus^strong^) progenitor state to the Notch-attenuated (Venus^weak^) redifferentiation state, leading to defective hepatocyte maturation during liver regeneration.

### *irf6* loss selectively reduces NF-*κ*B-active regenerating hepatocytes at A36h-R24h without altering progenitor-associated NF-*κ*B or Wnt signaling

Because biliary-driven liver regeneration is modulated by microenvironmental and pro-survival pathways—most notably NF-κB and canonical Wnt/beta-catenin signaling [12, 18]—we next investigated whether the impaired hepatocyte redifferentiation and reduced liver regrowth in *irf6* mutants are associated with alterations in these pathways. Using the *Tg(fabp10a:*CFP-NTR*);Tg(Tp1:*H2BmCherry) background crossed with either the NF-κB reporter *Tg(NF-κB:*EGFP) or the canonical Wnt/beta-catenin reporter *Tg(Tcf/Lef:*dGFP), we first verified that under unablated homeostatic conditions at 5 and 6 dpf, the numbers of EGFP^+^ NF-κB-responsive cells and dGFP^+^ Wnt-responsive cells within both the CFP^+^ hepatocyte and mCherry^+^ biliary compartments were comparable between *irf6^+/-^*control and *irf6^-/-^* mutant larvae (Supplementary Fig. 6). Basal dGFP^+^ Wnt/beta-catenin reporter activity remained minimal in both genotypes (Supplementary Fig. 6C, D). Following Mtz-induced hepatocyte ablation (Fig. 7A and Supplementary Fig. 7A), we tracked Wnt/beta-catenin and NF-κB reporter dynamics across late ablation (A33h) and regeneration stages (A36h-R6h and A36h-R24h). Analysis of *Tg(Tcf/Lef:*dGFP) larvae revealed no significant differences in the numbers of dGFP^+^ or dGFP^-^cells within the mCherry^+^ population at A33h and A36h-R6h, nor within the CFP^+^, mCherry^strong+^, or mCherry^weak+^ populations at A36h-R24h between *irf6^+/-^* and *irf6^-/-^* larvae (Supplementary Fig. 7B-D), indicating that canonical Wnt/beta-catenin activation proceeds independently of *irf6*. Similarly, analysis of *Tg(NF-κB:*EGFP) larvae at A33h and A36h-R6h showed that while EGFP^+^ cells were actively recruited to the regenerating liver region, the numbers of EGFP^+^ and EGFP^-^cells within the mCherry^+^ Notch-responsive LPC population did not differ significantly between *irf6^+/-^* control and *irf6^-/-^* mutant larvae (Fig. 7B, C). At A36h-R24h, when newly redifferentiated CFP^+^ hepatocytes actively emerged from mCherry^+^ LPCs to restore hepatic mass, we further stratified reporter activity across hepatocyte CFP^+^ and biliary-derived mCherry^strong+^ and mCherry^weak+^ compartments (Fig. 7D). Notably, while no significant differences were observed in the EGFP^+^ or EGFP^-^fractions within either the mCherry^strong+^ or mCherry^weak+^ populations, the number of CFP^+^/EGFP^+^ cells (NF-κB reporter-positive regenerating hepatocytes) was significantly reduced in *irf6^-/-^* mutant larvae compared with *irf6^+/-^* controls (Fig. 7D). Collectively, these results demonstrate that *irf6* loss does not perturb NF-κB or Wnt/beta-catenin signaling within Tp1-positive LPCs during early regeneration, but leads to a specific reduction in NF-κB-active regenerating hepatocytes at A36h-R24h, consistent with impaired LPC-to-hepatocyte redifferentiation and the attenuated hepatic mass recovery observed in *irf6* mutants.

**Fig. 7.**
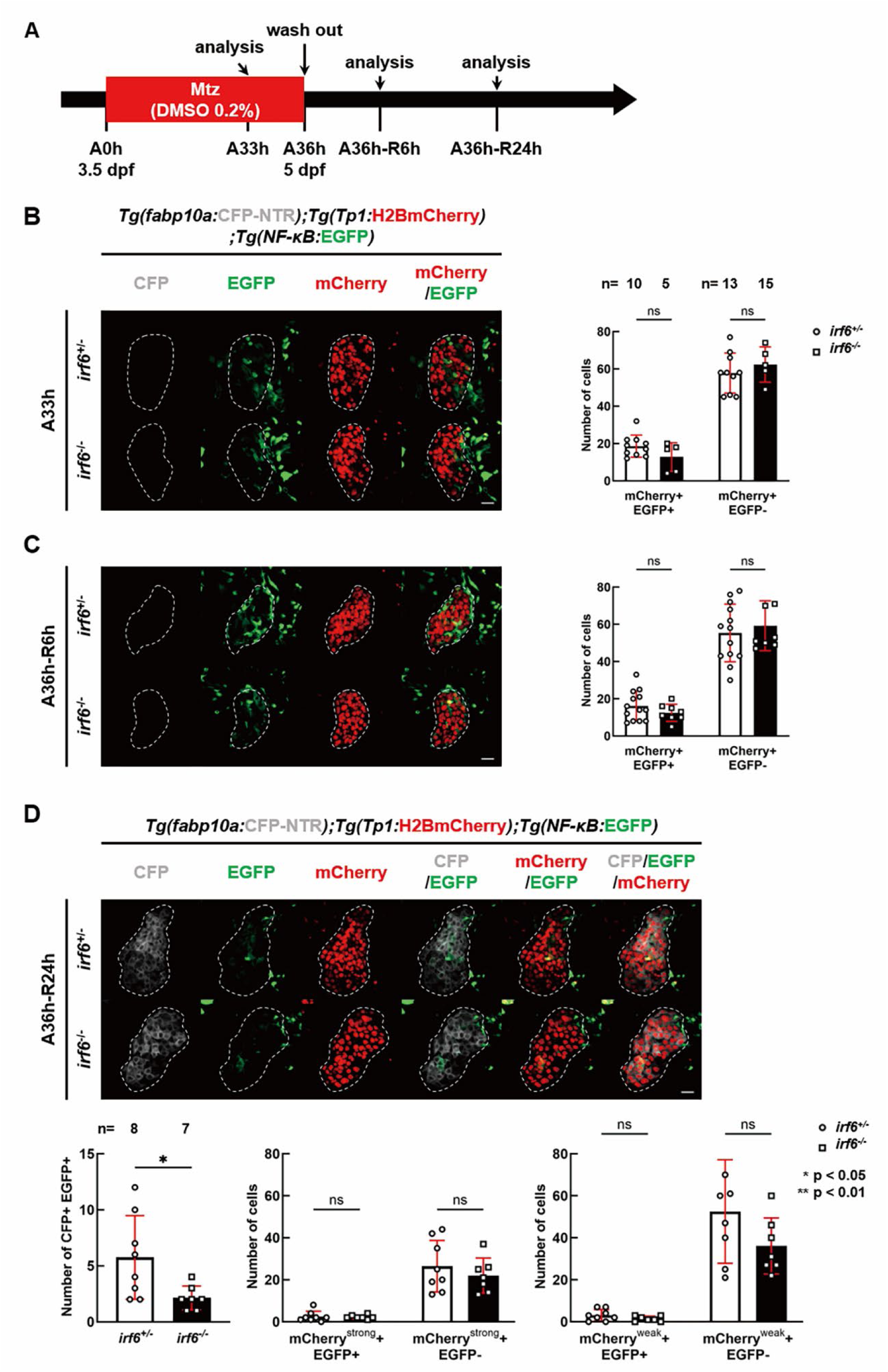
NF-*κ*B reporter activity during liver regeneration in *irf6* mutant larvae. (A) Experimental scheme illustrating Mtz-induced (in 0.2% DMSO) hepatocyte ablation from 3.5 to 5 dpf (A0h to A36h), followed by Mtz washout and confocal analysis at A33h, A36h-R6h, and A36h-R24h. (B–D) Representative confocal projection images (left/top) and quantitative analyses (right/bottom) of the regenerating liver region in *irf6^+/-^* control and *irf6^-/-^* mutant larvae at A33h (B), A36h-R6h (C), and A36h-R24h (D) using the *Tg(fabp10a:*CFP-NTR*);Tg(Tp1:*H2BmCherry*);Tg(*NF-κB:EGFP) reporter line. CFP (gray) labels hepatocytes, EGFP (green) labels NF-κB reporter-positive cells, and H2BmCherry (red) labels Tp1-positive Notch-responsive biliary-derived cells. Quantification at A33h (B) and A36h-R6h (C) showed no significant differences in the numbers of EGFP^+^ or EGFP^-^cells within the mCherry^+^ population between *irf6^+/-^* (white bars, circles) and *irf6^-/-^* (black bars, squares) larvae. At A36h-R24h (D), the number of CFP^+^/EGFP^+^ hepatocytes was significantly reduced in *irf6^-/-^* mutants compared with *irf6^+/-^*controls (bottom left), whereas EGFP^+^ and EGFP^-^cell numbers within the mCherry^strong+^ (bottom middle) and mCherry^weak+^ (bottom right) populations showed no significant differences. Each dot represents an individual larva. Red bars indicate the mean **±** SD. Statistical significance was determined using an unpaired two-tailed Student’s t test (\**P < 0.05*; \*\**P < 0.01*; \*\*\**P < 0.001*; ns., not significant). Scale bars: 20 μm.

## Discussion

In this study, we identify a previously unrecognized role for *irf6* in regulating hepatocyte redifferentiation during LPC-driven liver regeneration. *irf6* has been extensively characterized as a core transcription factor required for epithelial differentiation, periderm formation, and tissue morphogenesis during embryonic development [1–3, 6]; however, its function in post-embryonic tissue contexts, particularly during organ regeneration, has remained largely unexplored. Our developmental analyses confirmed that *irf6* is broadly expressed during early zebrafish embryogenesis and progressively becomes restricted to specific tissues, including endoderm-derived organs such as the liver (Fig. 1). Consistent with the results of previous studies [7, 8], maternal-zygotic loss of *irf6* (mz-*irf6^wku2001^*) resulted in severe early epiboly arrest and superficial epithelial disintegration, demonstrating an essential requirement for maternally supplied *irf6* during early embryogenesis (Fig. 2G and Supplementary Video 1). In contrast, zygotic *irf6* mutants exhibited normal primary liver development and hepatocyte differentiation under homeostatic conditions, indicating that zygotic *irf6* is dispensable for initial liver organogenesis (Fig. 2A–F).

Following targeted hepatocyte ablation, loss of zygotic *irf6* significantly impaired regenerative liver growth across recovery stages (Fig. 3 and Supplementary Fig. 1). Liver regeneration after severe parenchymal injury requires precise coordination of epithelial cell plasticity, proliferation, and redifferentiation [10]. While regeneration under mild injury conditions is largely driven by the self-duplication of pre-existing hepatocytes [11, 12], near-total hepatocyte loss activates an alternative regenerative program in which BECs dedifferentiate into proliferative hepatoblast-like LPCs that subsequently redifferentiate into mature hepatocytes [13–15]. Concordantly, we found that *irf6* transcription, which was largely quiescent in the homeostatic larval liver by 5–7 dpf, was actively re-induced and sustained within the regenerating hepatic niche throughout late ablation and intermediate regeneration windows (Fig. 4). Whole-mount expression profiling further revealed that key sequential regulators of hepatic competence and specification (*foxa3* and *hhex*), progenitor maintenance and differentiation transcription factors (*prox1* and *hnf4α*), and mature hepatocyte markers (*lfabp*, *gc*, *cp*, and *bhmt*) were qualitatively preserved in *irf6* mutants (Supplementary Fig. 2 and Supplementary Fig. 3), indicating that *irf6* deficiency does not cause a global loss of hepatic lineage identity, but rather compromises the quantitative efficiency of cellular transitions during LPC-driven liver regeneration.

A central finding of this study is that *irf6* is not required for the initial generation of Tp1-positive Notch-responsive biliary-derived LPCs, but is essential for their subsequent transition into redifferentiating hepatocytes (Fig. 5, Fig. 6, Supplementary Fig. 4, and Supplementary Fig. 5). Notch signaling plays a dual, stage-specific role during biliary-driven liver regeneration: high Notch activity is required for initial BEC activation and LPC maintenance, whereas subsequent Notch signal attenuation is strictly required for LPCs to exit the progenitor state and redifferentiate into functional hepatocytes [14, 17, 23, 24]. Using the dual-fluorescent *Tg(Tp1:*VenusPEST*);Tg(Tp1:*H2BmCherry) reporter system—in which high-Notch progenitors retain strong Venus fluorescence (Venus^strong^/mCherry^+^) whereas cells undergoing hepatocyte redifferentiation downregulate Notch to become Venus^weak^/mCherry^+^—we demonstrated that the total pool of Tp1-positive cells and Prox1^+^ progenitors was largely preserved in *irf6* mutants (Fig. 5B, Supplementary Fig. 4B). Instead, loss of *irf6* caused transitioning cells expressing the hepatoblast/hepatocyte transcription factor Hnf4α and the mature hepatocyte marker Bhmt to abnormally accumulate within the Notch-active Venus^strong^/mCherry^+^ compartment, accompanied by a deficit in transitioning into the Notch-attenuated Venus^weak^/mCherry^+^ redifferentiation state (Fig. 5C, Fig. 6B, Supplementary Fig. 4C, Supplementary Fig. 5B). Thus, *irf6* acts at a critical differentiation checkpoint downstream of LPC activation, promoting the timely attenuation of Notch signaling required for terminal hepatocyte redifferentiation.

Mechanistically, our findings align with and extend known regulatory links between IRF6, Notch signaling, and epithelial differentiation established in other developmental systems [1, 5, 6, 27, 28]. In mammalian keratinocytes, Notch1 directly activates IRF6 transcription to drive the switch from proliferation to terminal differentiation [27, 28], while IRF6 reciprocally promotes the proteasomal degradation of the progenitor-maintenance factor DeltaNp63 and activates downstream epithelial differentiation effectors such as Grhl3 [1, 6, 29]. By analogy, during LPC-driven liver regeneration, high Notch activity in activated BECs may initially induce *irf6* expression within the regenerating liver bud; once accumulated, Irf6 may function within a negative feedback loop to dampen Notch transcriptional output (facilitating the Venus^strong^-to-Venus^weak^ transition) and cooperate with lineage regulators such as Hnf4α to re-establish mature hepatocyte architecture. In our model, in the absence of *irf6*, transitioning LPCs initiate Hnf4α and Bhmt expression while retaining high Notch activity, trapping cells in an unresolved intermediate state that limits functional parenchymal expansion.

To exclude the possibility that this redifferentiation defect is driven by alterations in other major regenerative signaling axes, we also evaluated inflammatory NF-κB and canonical Wnt/beta-catenin signaling (Fig. 7, Supplementary Fig. 6, Supplementary Fig. 7). In the immune system and injured epithelia, NF-κB serves as a central coordinator of innate immunity and tissue repair, orchestrating macrophage/Kupffer cell activation and inflammatory cytokine production while simultaneously providing essential pro-survival and mitogenic signals within regenerating hepatocytes [20–22]. Previous immunological and epithelial studies have suggested that IRF6 can participate in Toll-like receptor–MyD88–IRAK4 cascades to modulate NF-κB activity either directly or indirectly depending on cellular context [30, 31]. In our *in vivo* lineage-resolved analyses, however, NF-κB reporter activity, *Tg(NF-κB:*EGFP), within Tp1-positive (mCherry^+^) biliary-derived progenitors—including both mCherry^strong+^ and mCherry^weak+^ subpopulations—remained completely unaltered in *irf6* mutants across all examined stages (A33h, A36h-R6h, and A36h-R24h; Fig. 7B–D). Instead, *irf6^-/-^* mutants exhibited a selective reduction only in CFP^+^/EGFP^+^ regenerating hepatocytes at A36h-R24h (Fig. 7D). This compartment-specific pattern indicates that *irf6* does not directly regulate NF-κB activation within Notch-responsive LPCs or the local inflammatory microenvironment during early regeneration. Rather, the reduction in NF-κB-positive hepatocytes at A36h-R24h represents an indirect, secondary consequence of impaired LPC-to-hepatocyte redifferentiation: because fewer Tp1-positive progenitors successfully attenuate Notch signaling to mature into CFP^+^ hepatocytes in *irf6* mutants, the pool of newly generated hepatocytes engaging pro-survival NF-κB signaling during parenchymal expansion is secondarily diminished. Furthermore, although canonical Wnt/beta-catenin signaling is a well-established driver of hepatocyte proliferation and LPC-to-hepatocyte specification following severe liver injury [14, 19], our analyses using the *Tg(Tcf/Lef:*dGFP) reporter showed no differences in Wnt/beta-catenin activity within either mCherry^+^ LPCs or CFP^+^ hepatocytes between control and *irf6^-/-^* larvae under basal (5 and 6 dpf; Supplementary Fig. 6C, D) or regenerating conditions (A33h to A36h-R24h; Supplementary Fig. 7). This demonstrates that *irf6* operates independently of canonical Wnt/beta-catenin signaling during liver regeneration. Thus, while extrinsic pathways such as Wnt/beta-catenin and Notch initiate progenitor mobilization and lineage specification [17, 19], intrinsic epithelial transcription factors such as *irf6* are specifically required to resolve progenitor signaling states—specifically by promoting Notch attenuation—to complete terminal differentiation.

Although our lineage-resolved analyses demonstrated that NF-κB activity within Tp1-positive LPCs remained intact in *irf6^-/-^* mutants, we observed robust recruitment of EGFP^+^ non-biliary cells around the regenerating liver niche during late ablation and early regeneration (Fig. 7B, C). Following severe hepatocyte ablation, infiltrating macrophages and resident Kupffer cells play critical roles in clearing necrotic cellular debris, remodeling the extracellular matrix, and secreting paracrine factors that support LPC expansion and hepatocyte maturation [20–22]. Given that IRF6 participates in innate immune Toll-like receptor (TLR) signaling cascades [30, 31] and that our *irf6^wku2001^* model represents a global loss-of-function allele, it remains possible that *irf6* also influences macrophage dynamics or epithelial–macrophage crosstalk within the regenerative microenvironment. Future studies utilizing macrophage-specific transgenic reporters, targeted macrophage depletion, and cell-type-specific rescue approaches will be valuable to determine whether *irf6* directly modulates macrophage recruitment, phagocytic clearance, or phenotypic polarization, and how macrophage-derived signals coordinate with *irf6*-mediated Notch attenuation to drive efficient hepatocyte redifferentiation.

In summary, using a zebrafish model of severe hepatocyte ablation, we demonstrated that *irf6* is a key intrinsic regulator of hepatocyte redifferentiation efficiency during LPC-driven liver regeneration. Loss of *irf6* impairs regenerative liver outgrowth, disrupts the transition of Tp1-positive progenitors from a Notch-active (Venus^strong^) state to a Notch-attenuated (Venus^weak^) redifferentiation state, and secondarily reduces the emergence of NF-κB-active regenerating hepatocytes, without directly altering progenitor-associated NF-κB or Wnt/beta-catenin signaling. These findings expand the functional repertoire of *irf6* beyond embryonic epithelial morphogenesis and provide new insight into the transcriptional mechanisms that govern cell-state transitions during organ regeneration *in vivo*.

## Methods

### Zebrafish strains and maintenance

Zebrafish (*Danio rerio*) embryos and adult fish were raised and maintained under standard laboratory conditions. The *irf6^wku2001^*mutant allele was generated using CRISPR/Cas9-mediated genome editing following established zebrafish gene-targeting protocols. Single-guide RNAs (sgRNAs: 5′-GAAGCCCTGGCTGGTCTCTCAGG-3′, 5′-TGGACAACGCTACGTTTCCTGGG-3′) were designed to target the coding region of the *irf6* gene, inducing a 23-bp deletion and frameshift mutation that introduced a premature stop codon and produced a truncated Irf6 peptide (22 amino acids). The sgRNAs and Cas9 mRNA were co-injected into one-cell stage embryos, and founder fish carrying the mutation were identified and outcrossed to establish a stable mutant line. Genotyping of *irf6^wku2001^*mutants was performed via PCR amplification of genomic DNA using gene-specific primers (forward: 5′-TGTACAGCCGGTGTGATGTC-3′; reverse: 5′-GGTGTTCTCCTCCTCCTGCTG-3′). PCR was conducted under standard thermal cycling conditions (95 °C for 5 min; 35 cycles of 95 °C for 30 s, 60 °C for 30 s, and 72 °C for 10 s; followed by a final extension at 72 °C for 5 min), and PCR amplicons were resolved using agarose gel electrophoresis. Time-lapse imaging of early maternal-zygotic (mz-*irf6^wku2001^*) mutant and wild-type embryos was performed from cleavage through gastrulation stages using a stereomicroscope. The mutant and transgenic lines were as follows: *irf6^wku2001^*, *Tg(fabp10a:CFP-NTR)^s931^*, *Tg(fabp10a:DsRed)^gz15^*, *Tg(EPV.Tp1-Mmu.Hbb:hist2h2l-mCherry)^s939^*, *Tg(EPV.Tp1-Mmu.Hbb:Venus-Mmu.Odc1)^s940^*, *Tg(6xHsa.NFκB:EGFP)^nc1^*, and *Tg(Tcf/Lef-miniP:dGFP)^kyu1^* [referred to as *Tg(fabp10a:CFP-NTR)*, *Tg(fabp10a:DsRed)*, *Tg(Tp1:H2BmCherry)*, *Tg(Tp1:VenusPEST)*, *Tg(NF-κB:EGFP),* and *Tg(Tcf/Lef:dGFP)*, respectively].

### Reverse-transcription PCR analysis

To analyze the temporal expression of *irf6* during zebrafish development, embryos and larvae were collected at the indicated developmental stages ranging from the two-cell stage to 5 days post-fertilization (dpf). Total RNA was extracted from pooled samples at each stage, followed by DNase treatment and cDNA synthesis via reverse transcription. Conventional reverse-transcription (RT)-PCR was performed using gene-specific primers for *irf6*, with *β-actin* used as an internal loading control. PCR products were resolved using agarose gel electrophoresis and visualized to assess relative transcript levels across developmental stages.

### Hepatocyte ablation

Targeted hepatocyte ablation was performed using the NTR/Mtz system. Larvae carrying the *Tg(fabp10a:CFP-NTR)* transgene were treated with 10 mM metronidazole (Mtz; Sigma-Aldrich, St. Louis, MO, USA) dissolved in egg water supplemented with 0.2% dimethyl sulfoxide (DMSO) and 0.2 mM 1-phenyl-2-thiourea (Sigma-Aldrich) for 36 hours from 3.5 dpf (ablation 0 h, A0h) to 5 dpf (A36h) to induce hepatocyte-specific ablation [13, 25]. Following ablation at A36h, Mtz was washed out using fresh egg water, and larvae were allowed to recover until the indicated regeneration time points (A36h-R6h, A36h-R24h, A36h-R30h, A36h-R48h, and A36h-R72h).

### Whole-mount *in situ* hybridization and immunofluorescence analysis

Whole-mount *in situ* hybridization (WISH) was performed using digoxigenin-labeled antisense RNA probes following standard protocols. Antisense RNA probes were synthesized for *irf6*, mature hepatocyte markers (*bhmt*, *cp*, *lfabp*, and *gc*) [26, 32, 33], and hepatic progenitor- and lineage-associated transcription factors (*hhex*, *foxa3*, *hnf4α*, and *prox1*) [14, 26, 34]. After staining, embryos and larvae were cleared and imaged using a Stemi stereo microscope (Carl Zeiss, Oberkochen, Germany). Whole-mount immunofluorescence was performed on *Tg(fabp10a:CFP-NTR);Tg(Tp1:VenusPEST);Tg(Tp1:H2BmCherry)* reporter larvae using the following primary antibodies: rabbit anti-Bhmt (1:100; GeneTex, Inc., Irvine, CA, USA), rabbit anti-Hnf4α (1:20; Sigma-Aldrich), and rabbit anti-Prox1 (1:100; GeneTex, Inc., Irvine, CA, USA), followed by Alexa Fluor 647-conjugated secondary antibodies (1:200; Invitrogen, Carlsbad, CA, USA).

### Confocal imaging

Fluorescence and immunofluorescence images were acquired using an LSM 980 confocal microscope (Carl Zeiss) at the Core Facility for Supporting Analysis & Imaging of Biomedical Materials at Wonkwang University, supported by the National Research Facilities and Equipment Center. For Tp1 Notch reporter-based lineage analysis, VenusPEST-positive and H2BmCherry-positive cells were identified in *Tg(fabp10a:CFP-NTR);Tg(Tp1:VenusPEST);Tg(Tp1:H2BmCherry)* larvae. Venus^+^/mCherry^+^ cells were quantified as Tp1-positive Notch-responsive biliary-derived cells and further stratified based on VenusPEST fluorescence intensity into high-Notch (Venus^strong^/mCherry^+^) and Notch-attenuated (Venus^weak^/mCherry^+^) subpopulations. Cells positive for Prox1, Hnf4α, or Bhmt together with Venus and H2BmCherry were quantified as marker-positive Tp1-derived cells within each reporter state. For NF-κB and Wnt/beta-catenin reporter analyses, larvae carrying *Tg(NF-κB:EGFP)* or *Tg(Tcf/Lef:dGFP)* in the *Tg(fabp10a:CFP-NTR);Tg(Tp1:H2BmCherry)* background were imaged at unablated developmental stages (5 and 6 dpf) and at defined time points following Mtz-induced hepatocyte ablation (A33h, A36h-R6h, and A36h-R24h). EGFP^+^ (NF-κB-responsive) and dGFP^+^ (Wnt/beta-catenin-responsive) cells were identified in the liver region and quantified based on their colocalization with CFP^+^ hepatocytes or mCherry^+^ (mCherry^strong+^ and mCherry^weak+^) biliary-derived cells. Identical imaging settings and quantification criteria were applied across control and *irf6* mutant larvae.

### Liver size measurement

Regenerating liver size was quantified by measuring the two-dimensional DsRed-positive fluorescent liver area in *Tg(fabp10a:CFP-NTR);Tg(fabp10a:DsRed)* larvae using ImageJ software (NIH, Bethesda, MD, USA). Measurements were performed on lateral fluorescence images across defined regeneration stages.

### Statistical analysis

Statistical analyses and graph generation were performed using GraphPad Prism 10 software (GraphPad Software, San Diego, CA, USA). Each data point represents an individual larva. Data are presented as the mean **±** standard deviation (SD). Differences between groups were tested using an unpaired two-tailed Student’s t test, and a P value < 0.05 was considered statistically significant.

## Data Availability

All data supporting the findings of this study are available within the article and its Supplementary Information files. Source data are provided with this paper. Additional raw imaging data and biological materials, including the *irf6^wku2001^* zebrafish mutant line generated in this study, are available from the corresponding author upon reasonable request.

## Animal Ethics

All animal procedures and experimental protocols were reviewed and approved by the Institutional Animal Care and Use Committee (IACUC) of Wonkwang University and were conducted in strict accordance with institutional and national guidelines for the care and use of laboratory animals.

## Author Contributions

N.K., S.L., and T.-Y.C. conceived and designed the study. N.K. and S.L. performed the experiments, maintained the zebrafish lines, and acquired the data. N.K., S.L., and T.-Y.C. analyzed and interpreted the data. N.K. and T.-Y.C. wrote and revised the manuscript. T.-Y.C. supervised the project and acquired funding. All authors read and approved the final manuscript.

---

**Supplementary Video 1. Time-lapse imaging of early embryonic development in *irf6^+/+^* and mz-*irf6^-/-^* embryos.** Representative bright-field time-lapse movie comparing wild-type (*irf6^+/+^*, left, n = 9) and maternal-zygotic *irf6^wku2001^* (mz-*irf6^-/-^*, right, n = 9) embryos recorded over 335 min starting from the sphere stage (10 min). While *irf6^+/+^* embryos progressed normally through 50% epiboly (135 min), shield (210 min), 70% epiboly (285 min), and 90% epiboly (335 min), mz-*irf6^-/-^* embryos remained arrested at the sphere stage and underwent progressive superficial epithelial rupture and blastoderm disintegration (indicated by white arrows) beginning at 210 min. Scale bars: 500 μm.

**Supplementary Fig. 1.**
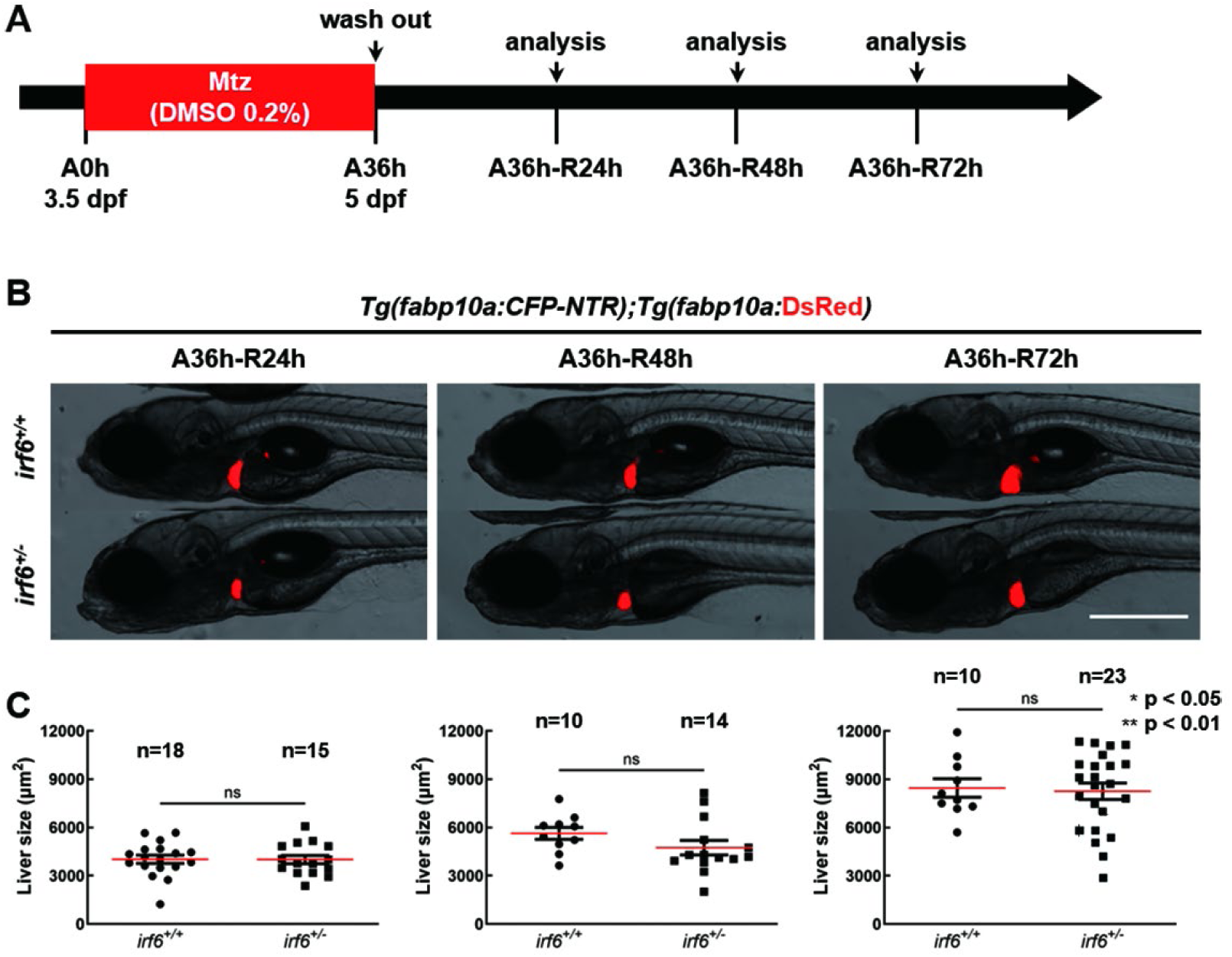
Comparable liver regeneration between wild-type (*irf6^+/+^*) and heterozygous (*irf6^+/-^*) larvae following hepatocyte ablation. (A) Experimental scheme showing the timeline of Mtz (in 0.2% DMSO) treatment for hepatocyte ablation from 3.5 to 5 dpf (A0h to A36h), followed by Mtz washout and subsequent analysis at A36h-R24h, A36h-R48h, and A36h-R72h. (B) Representative lateral fluorescence images merged with bright-field views showing regenerating livers (DsRed, red) at A36h-R24h, A36h-R48h, and A36h-R72h in *irf6^+/+^* and *irf6^+/-^* larvae carrying the *Tg(fabp10a:CFP-NTR);Tg(fabp10a:*DsRed) transgenes. (C) Quantification of regenerating liver size (μm^2^) comparing *irf6^+/+^* and *irf6^+/-^* larvae at A36h-R24h (left), A36h-R48h (middle), and A36h-R72h (right). Each data point represents an individual larva, and the number of examined larvae (n) is indicated above each group. Red horizontal bars indicate the mean, and black error bars indicate the standard deviation (SD). Statistical significance was assessed using an unpaired two-tailed Student’s t test (\**P < 0.05*; \*\**P < 0.01*; \*\*\**P < 0.001*; ns., not significant). Scale bar: 500 μm.

**Supplementary Fig. 2.**
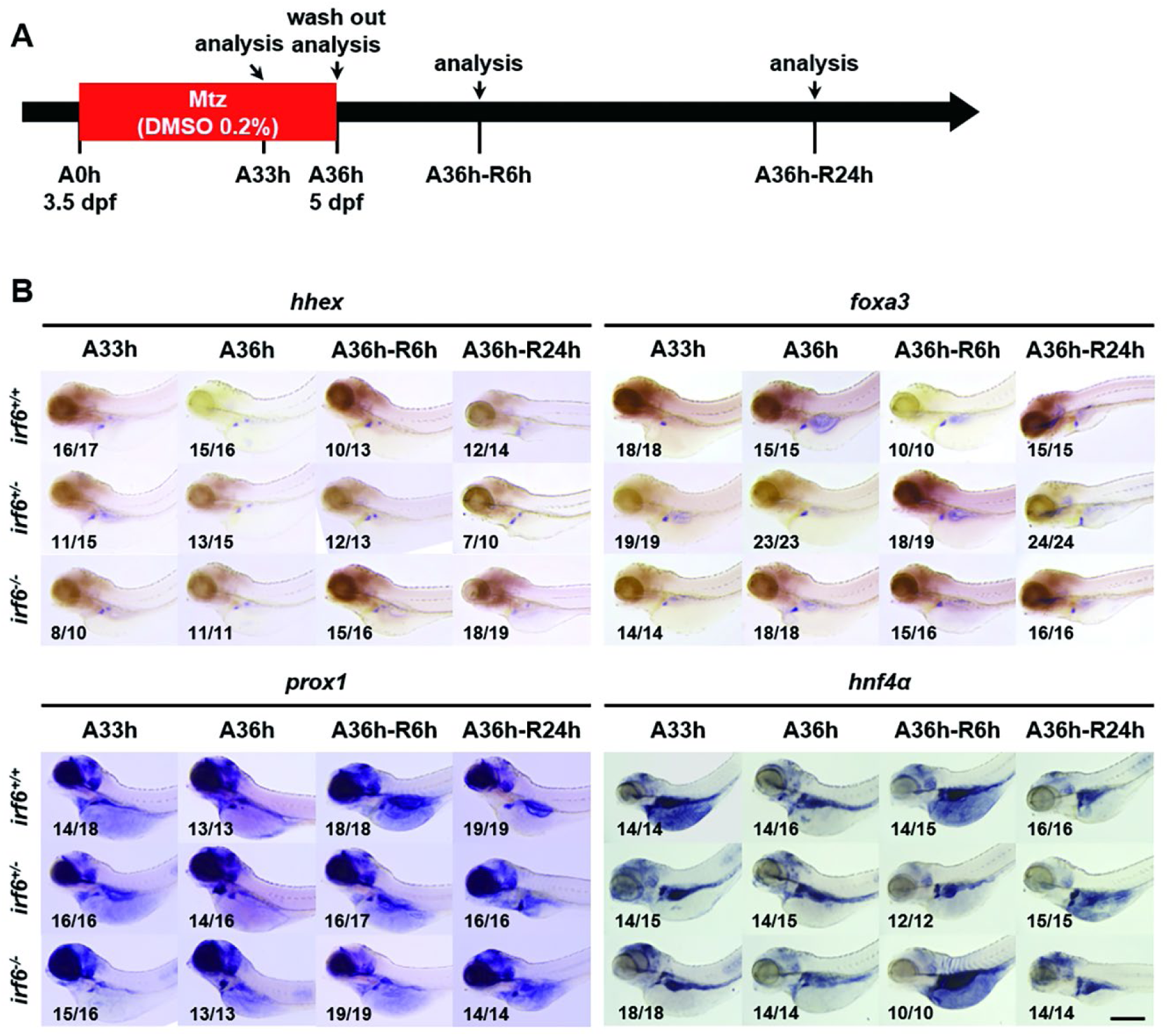
Expression patterns of hepatic competence, specification, and progenitor-associated transcription factors during early liver regeneration. (A) Experimental scheme illustrating Mtz-induced (in 0.2% DMSO) hepatocyte ablation from 3.5 to 5 dpf (A0h to A36h), followed by Mtz washout and WISH analysis at defined late-ablation and early regeneration stages (A33h, A36h, A36h-R6h, and A36h-R24h). (B) Representative lateral WISH images showing the expression of *hhex* and *foxa3* (sequential regulators of hepatic competence, specification, and liver bud morphogenesis) as well as *prox1* and *hnf4α* (cooperative yet contrasting regulators of hepatic progenitor maintenance and hepatocyte differentiation) in *irf6^+/+^*, *irf6^+/-^*, and *irf6^-/-^* larvae at A33h, A36h, A36h-R6h, and A36h-R24h. Numbers in the bottom-left corner of each panel indicate the number of larvae exhibiting the representative expression pattern out of the total number examined. Scale bar: 250 μm.

**Supplementary Fig. 3.**
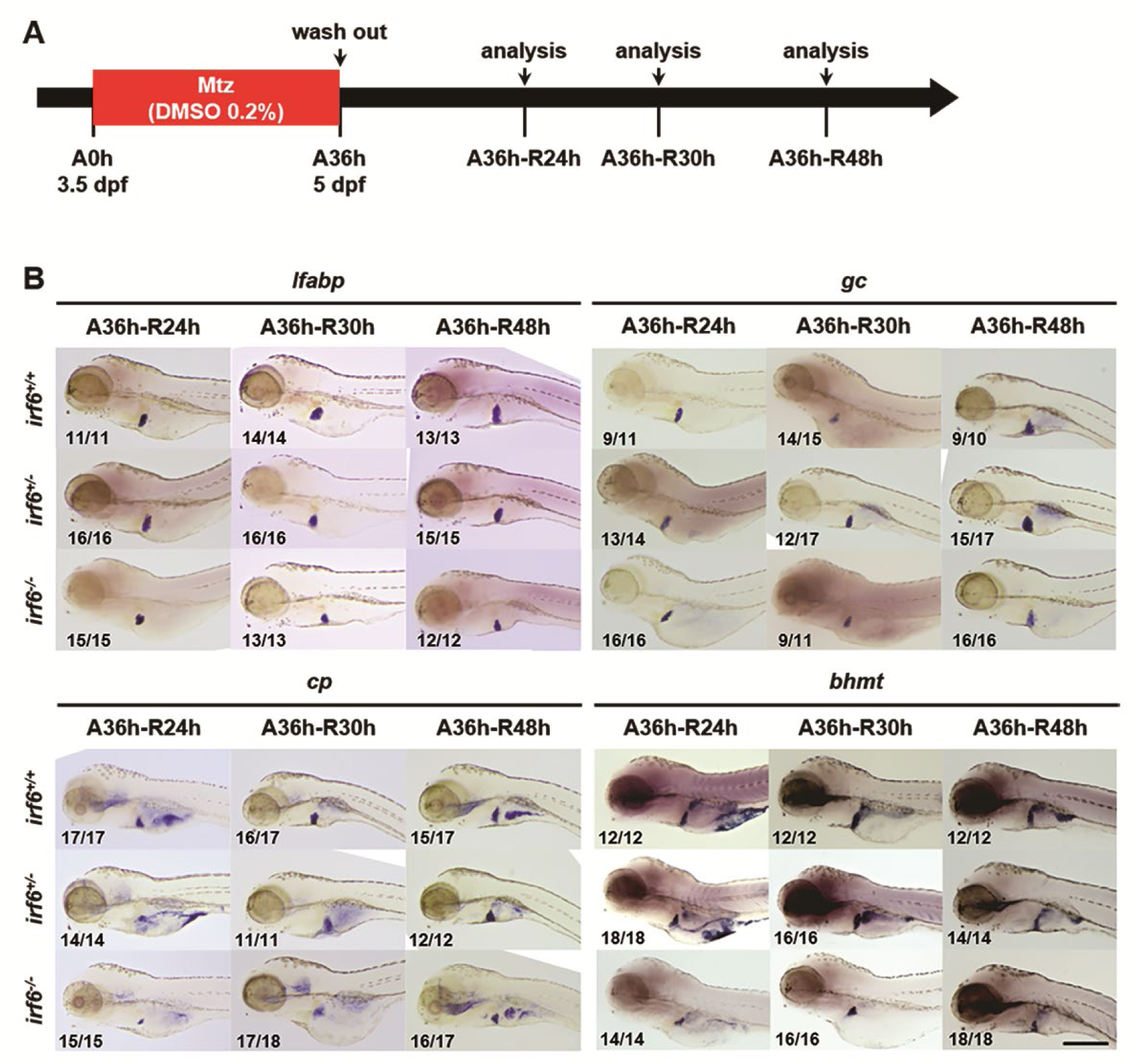
Expression patterns of mature hepatocyte markers during liver regeneration. (A) Experimental scheme illustrating Mtz-induced (in 0.2% DMSO) hepatocyte ablation from 3.5 to 5 dpf (A0h to A36h), followed by Mtz washout and WISH analysis at defined intermediate-to-late regeneration stages (A36h-R24h, A36h-R30h, and A36h-R48h). (B) Representative lateral WISH images showing the expression of mature hepatocyte markers (*lfabp*, *gc*, *cp*, and *bhmt*) in *irf6^+/+^*, *irf6^+/-^*, and *irf6^-/-^*larvae at A36h-R24h, A36h-R30h, and A36h-R48h. Numbers in the bottom-left corner of each panel indicate the number of larvae exhibiting the representative expression pattern out of the total number examined. Scale bar: 250 μm.

**Supplementary Fig. 4.**
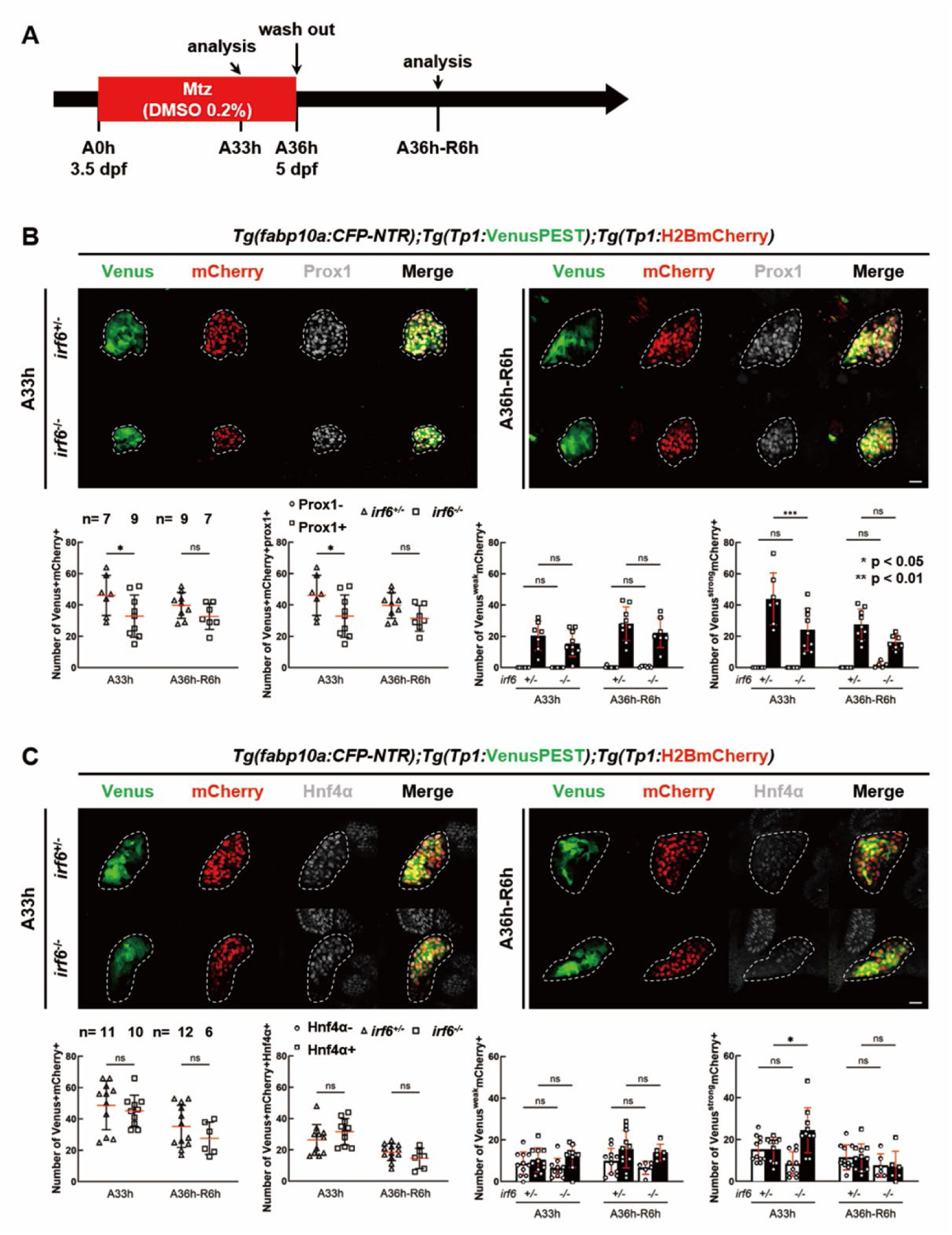
Absolute cell number quantification of Prox1^-^ and Hnf4*α*-expressing Tp1-positive populations during liver regeneration. (A) Experimental scheme illustrating Mtz-induced (in 0.2% DMSO) hepatocyte ablation from 3.5 to 5 dpf (A0h to A36h), followed by Mtz washout and analysis at A33h and A36h-R6h. (B) Representative confocal immunofluorescence images (top) and absolute cell number quantifications (bottom) of livers from *irf6^+/-^* and *irf6^-/-^* larvae at A33h and A36h-R6h in the *Tg(fabp10a:CFP-NTR);Tg(Tp1:*VenusPEST*);Tg(Tp1:*H2BmCherry) background immunostained for Prox1 (gray). Graphs show the total number of Venus^+^/mCherry^+^ cells (first), the number of Prox1^+^/Venus^+^/mCherry^+^ cells (second), and the numbers of Venus^weak^/mCherry^+^ (third) and Venus^strong^/mCherry^+^ (fourth) cells stratified by Prox1 expression (Prox1^-^, open circles/white bars; Prox1^+^, open squares/black bars). (C) Representative confocal immunofluorescence images (top) and absolute cell number quantifications (bottom) of livers from *irf6^+/-^* and *irf6^-/-^* larvae at A33h and A36h-R6h immunostained for Hnf4α (gray). Graphs show the total number of Venus^+^/mCherry^+^ cells (first), the number of Hnf4α^+^/Venus^+^/mCherry^+^ cells (second), and the numbers of Venus^weak^/mCherry^+^ (third) and Venus^strong^/mCherry^+^ (fourth) cells stratified by Hnf4α expression (Hnf4α^-^, open circles/white bars; Hnf4α^+^, open squares/black bars). Dashed lines outline the regenerating liver. Each data point represents an individual larva, with the number of examined larvae (n) indicated above the graphs. Red bars indicate the mean **±** SD. Statistical significance was determined using an unpaired two-tailed Student’s t test (\**P < 0.05*; \*\**P < 0.01*; \*\*\**P < 0.001*; ns., not significant). Scale bars: 20 μm.

**Supplementary Fig. 5.**
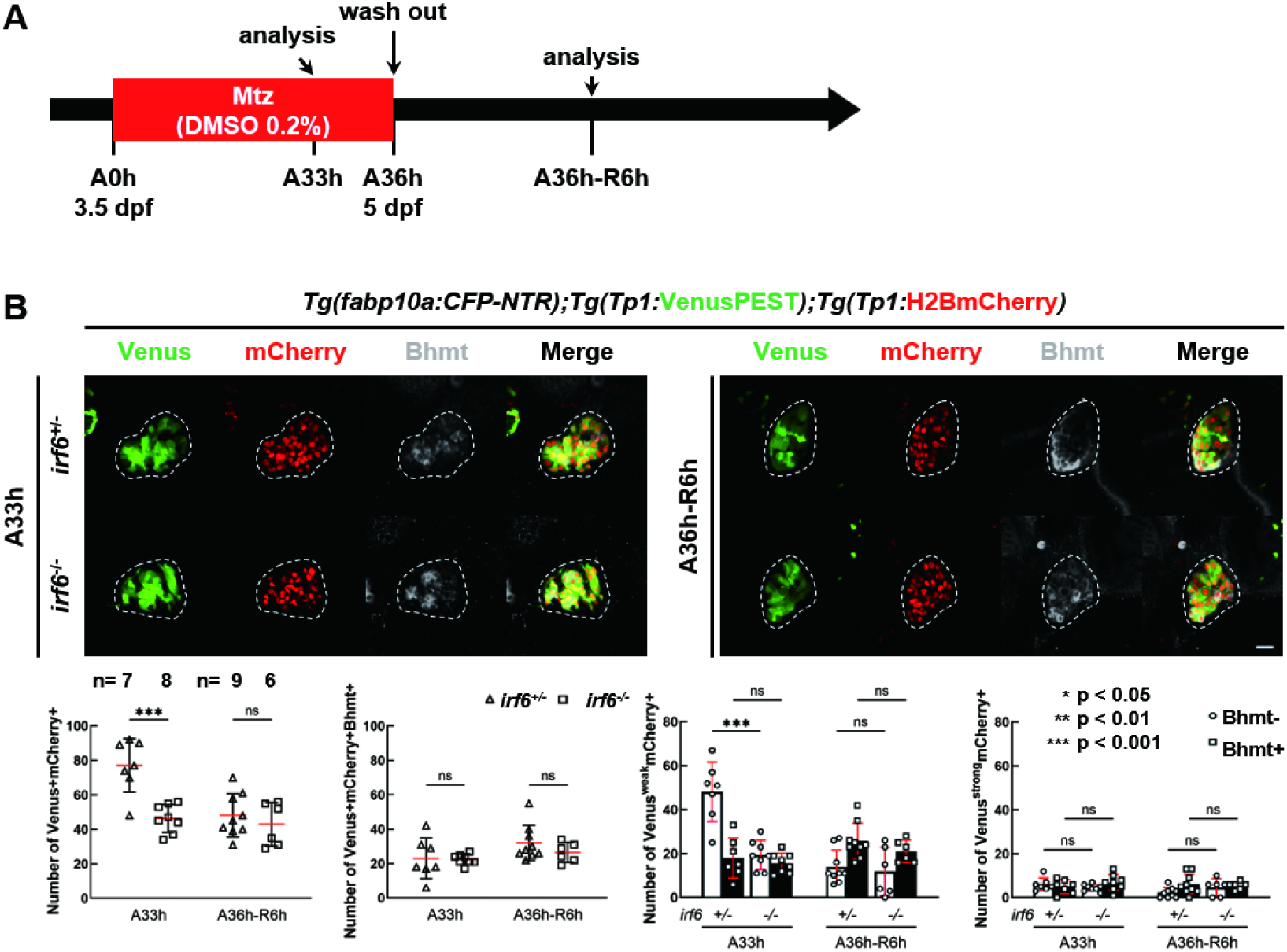
Absolute cell number quantification of Bhmt-expressing Tp1-positive populations during liver regeneration. (A) Experimental scheme illustrating Mtz-induced (in 0.2% DMSO) hepatocyte ablation from 3.5 to 5 dpf (A0h to A36h), followed by Mtz washout and immunofluorescence analysis at A33h and A36h-R6h. (B) Representative confocal immunofluorescence images (top) and absolute cell number quantifications (bottom) of livers from *irf6^+/-^* and *irf6^-/-^*larvae at A33h and A36h-R6h in the *Tg(fabp10a:CFP-NTR);Tg(Tp1:*VenusPEST*);Tg(Tp1:*H2BmCherry) background immunostained for Bhmt (gray). Graphs show the total number of Venus^+^/mCherry^+^ cells (first), the number of Bhmt^+^/Venus^+^/mCherry^+^ cells (second), and the numbers of Venus^weak^/mCherry^+^ (third) and Venus^strong^/mCherry^+^ (fourth) cells stratified by Bhmt expression (Bhmt^-^, open circles/white bars; Bhmt^+^, open squares/black bars). Dashed lines outline the regenerating liver. Each data point represents an individual larva, with the number of examined larvae (n) indicated above the graphs. Red bars indicate the mean **±** SD. Statistical significance was determined using an unpaired two-tailed Student’s t test (\**P < 0.05*; \*\**P < 0.01*; \*\*\**P < 0.001*; ns., not significant). Scale bar: 20 μm.

**Supplementary Fig. 6.**
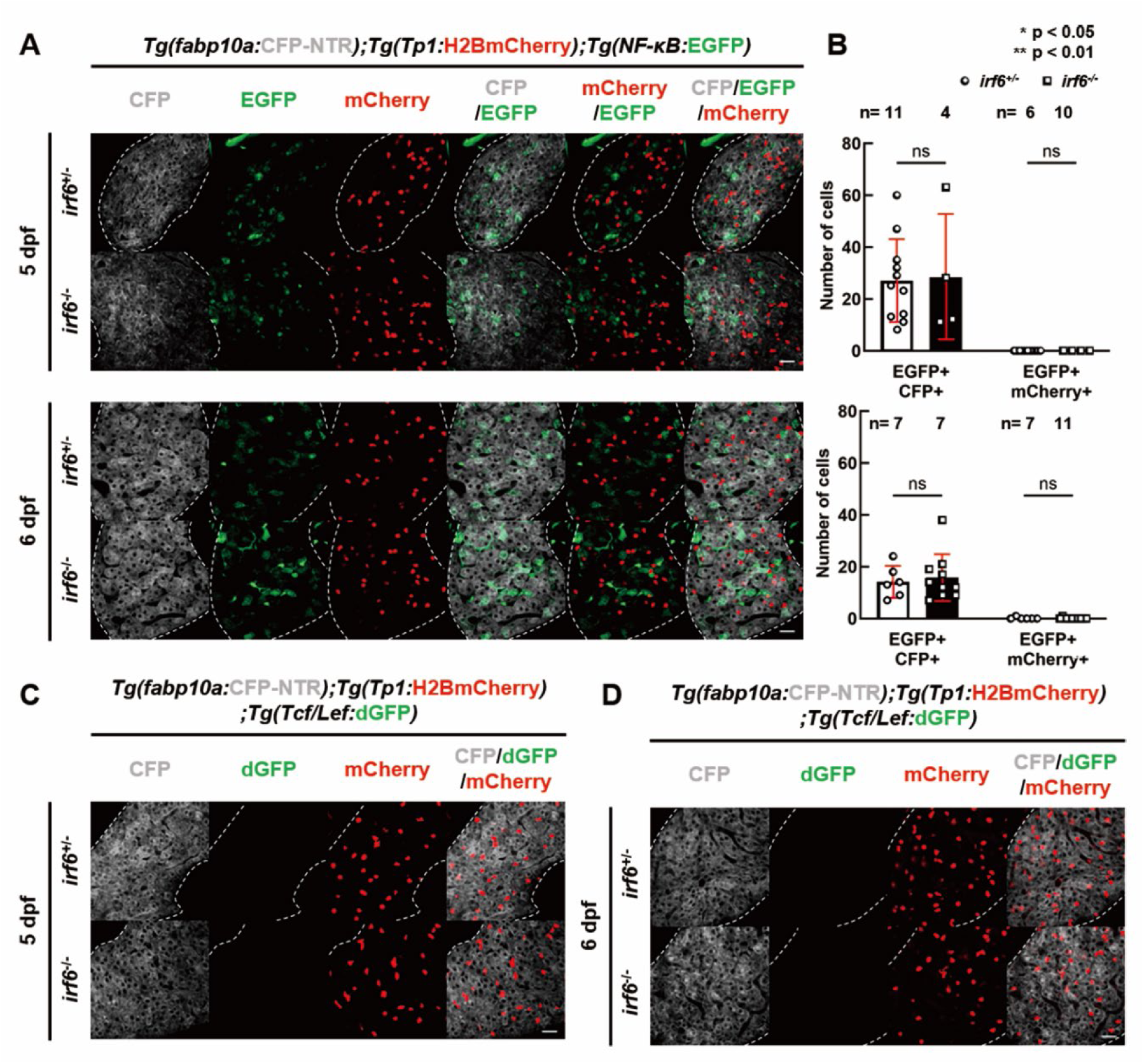
Basal NF-*κ*B and Wnt/beta-catenin reporter activities in the unablated larval liver at 5 and 6 dpf. (A, B) Representative confocal fluorescence images (A) and cell number quantifications (B) of unablated livers from *irf6^+/-^* control and *irf6^-/-^*mutant larvae at 5 dpf (top) and 6 dpf (bottom) carrying the *Tg(fabp10a:*CFP-NTR*);Tg(Tp1:*H2BmCherry*);Tg(NF-κB:*EGFP) transgenes. CFP (gray) labels hepatocytes, EGFP (green) labels NF-κB-responsive cells, and H2BmCherry (red) labels Tp1-positive biliary epithelial cells. Graphs show no significant differences in the numbers of EGFP^+^/CFP^+^ or EGFP^+^/mCherry^+^ cells between *irf6^+/-^* (open circles, white bars) and *irf6^-/-^* (open squares, black bars) larvae at either 5 or 6 dpf. (C, D) Representative confocal fluorescence images of unablated livers from *irf6^+/-^* and *irf6^-/-^* larvae at 5 dpf (C) and 6 dpf (D) carrying the *Tg(fabp10a:*CFP-NTR*);Tg(Tp1:*H2BmCherry*);Tg(Tcf/Lef:*dGFP) transgenes, which showed minimal basal dGFP (green, canonical Wnt/beta-catenin reporter) expression in both genotypes. Dashed lines outline the liver. Each data point represents an individual larva, with the number of examined larvae (n) indicated above the graphs. Red bars indicate the mean **±** SD. Statistical significance was determined using an unpaired two-tailed Student’s t test (\**P < 0.05*; \*\**P < 0.01*; \*\*\**P < 0.001*; ns., not significant). Scale bars: 20 μm.

**Supplementary Fig. 7.**
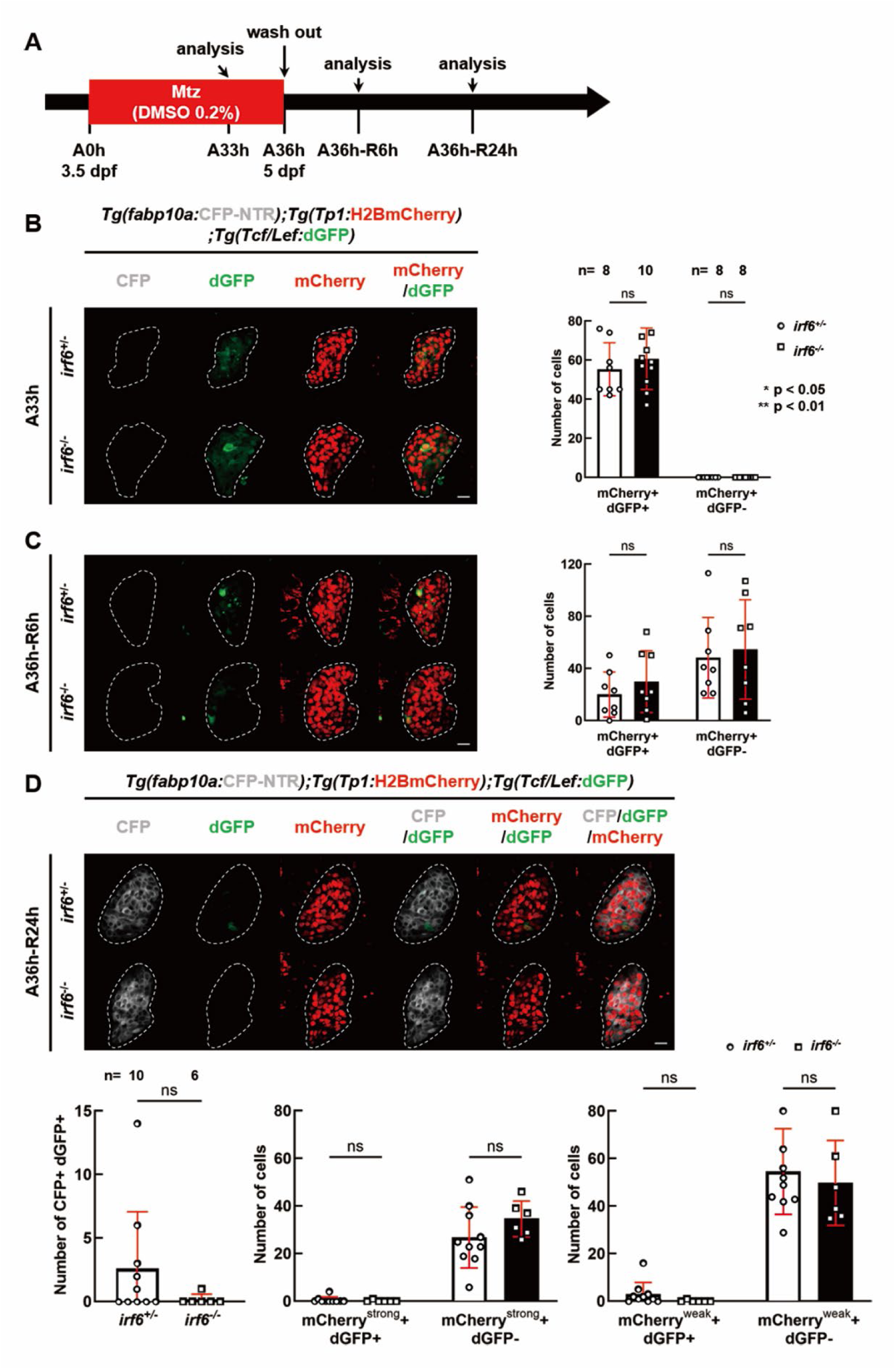
Canonical Wnt/beta-catenin reporter activity during liver regeneration in *irf6* mutant larvae. (A) Experimental scheme illustrating Mtz-induced (in 0.2% DMSO) hepatocyte ablation from 3.5 to 5 dpf (A0h to A36h), followed by Mtz washout and confocal analysis at A33h, A36h-R6h, and A36h-R24h. (B–D) Representative confocal fluorescence images and quantitative cell counts of the regenerating liver region in *irf6^+/-^* control and *irf6^-/-^* mutant larvae at A33h (B), A36h-R6h (C), and A36h-R24h (D) in the *Tg(fabp10a:*CFP-NTR*);Tg(Tp1:*H2BmCherry*);Tg(Tcf/Lef:*dGFP) background. CFP (gray) labels hepatocytes, dGFP (green) labels Wnt/beta-catenin reporter-positive cells, and H2BmCherry (red) labels Tp1-positive Notch-responsive cells. Quantifications at A33h (B) and A36h-R6h (C) showed no significant differences in the numbers of mCherry^+^/dGFP^+^ or mCherry^+^/dGFP^-^cells between *irf6^+/-^* (open circles, white bars) and *irf6^-/-^* (open squares, black bars) larvae. At A36h-R24h (D), no significant differences were observed in the number of CFP^+^/dGFP^+^ hepatocytes (bottom left) or in the numbers of dGFP^+^ and dGFP^-^cells within the mCherry^strong+^ (bottom middle) and mCherry^weak+^ (bottom right) populations. Dashed lines outline the regenerating liver. Each data point represents an individual larva, with the number of examined larvae (n) indicated above the graphs. Red bars indicate the mean **±** SD. Statistical significance was determined using an unpaired two-tailed Student’s t test (\**P < 0.05*; \*\**P < 0.01*; \*\*\**P < 0.001*; ns., not significant). Scale bars: 20 μm.

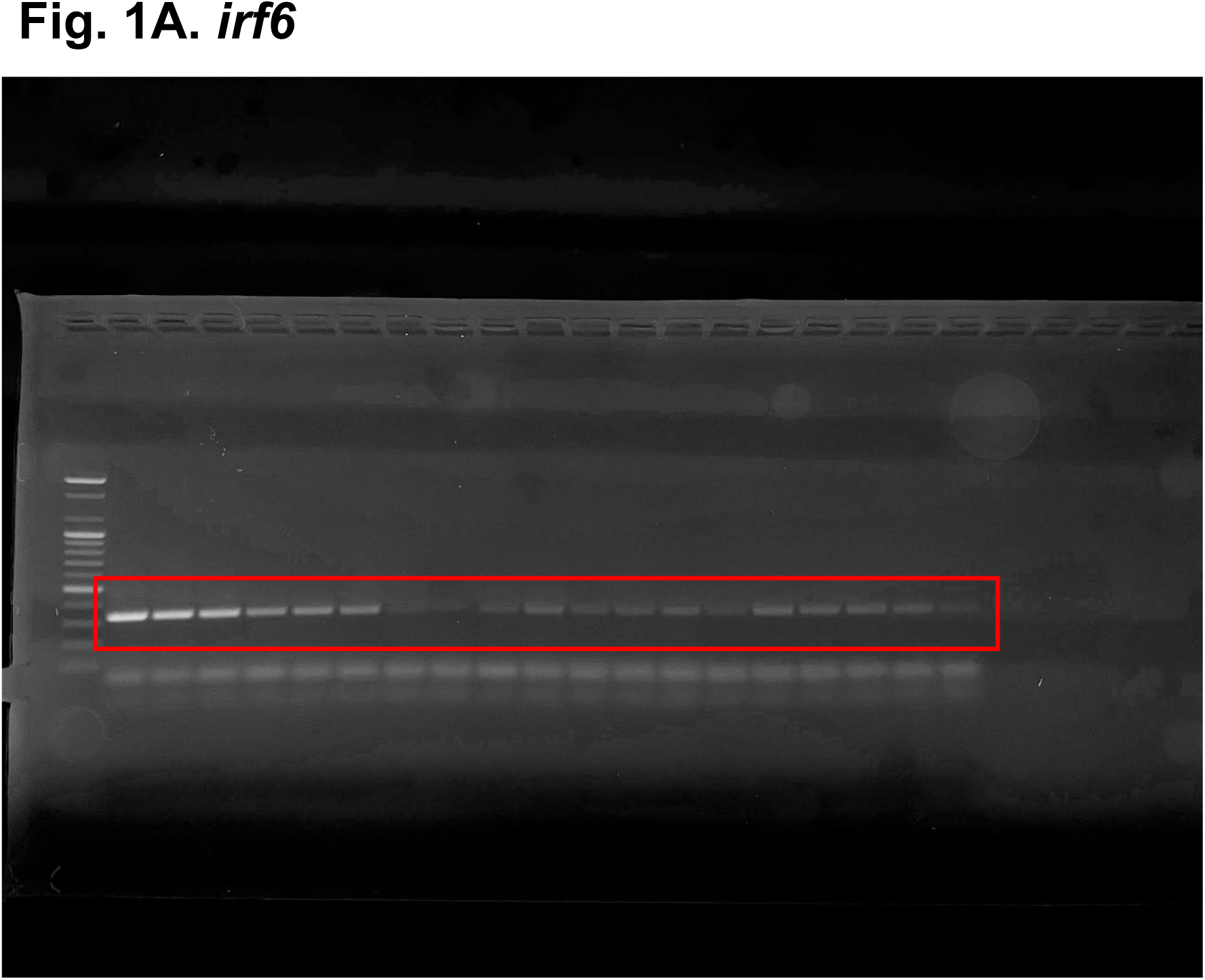

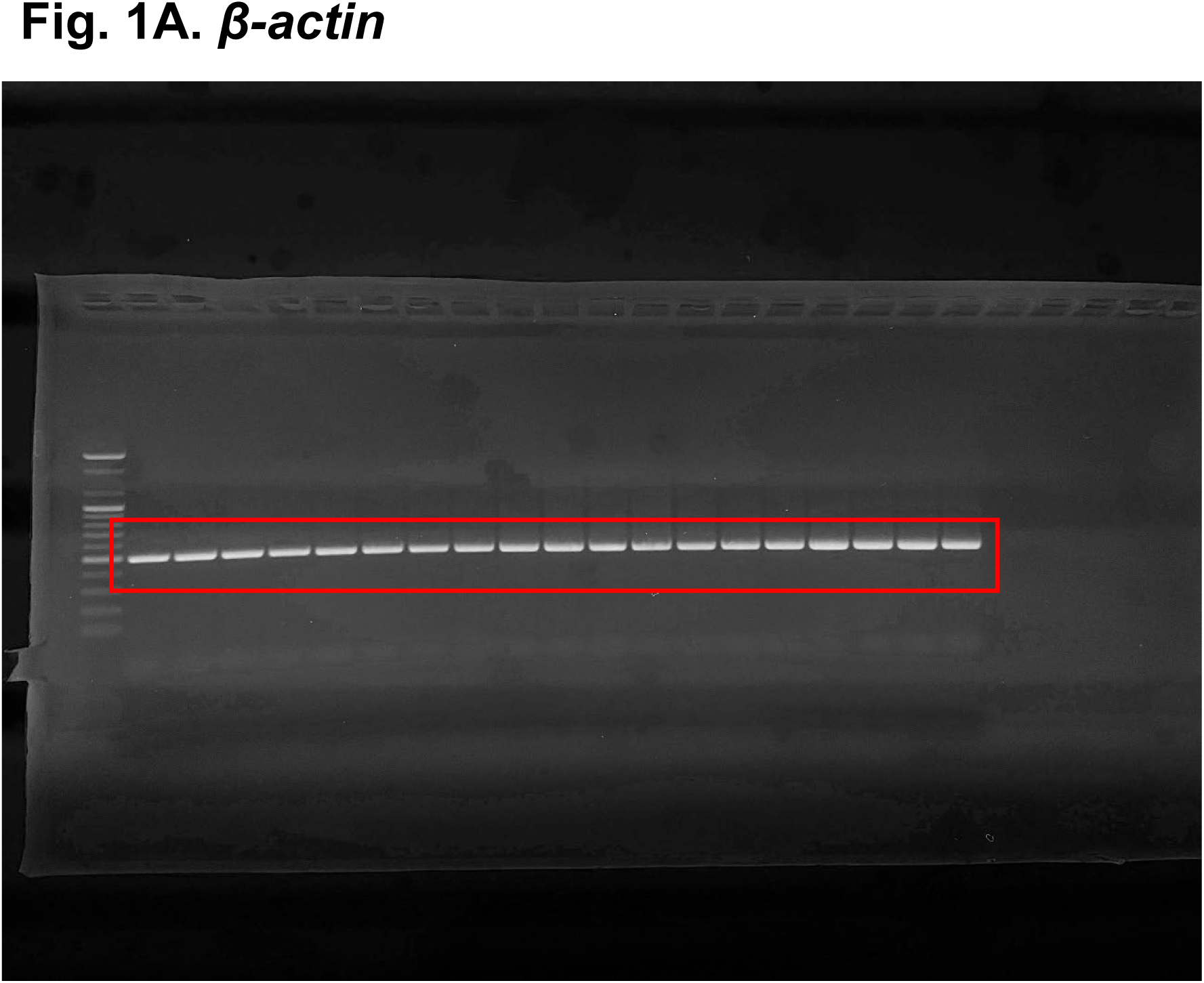

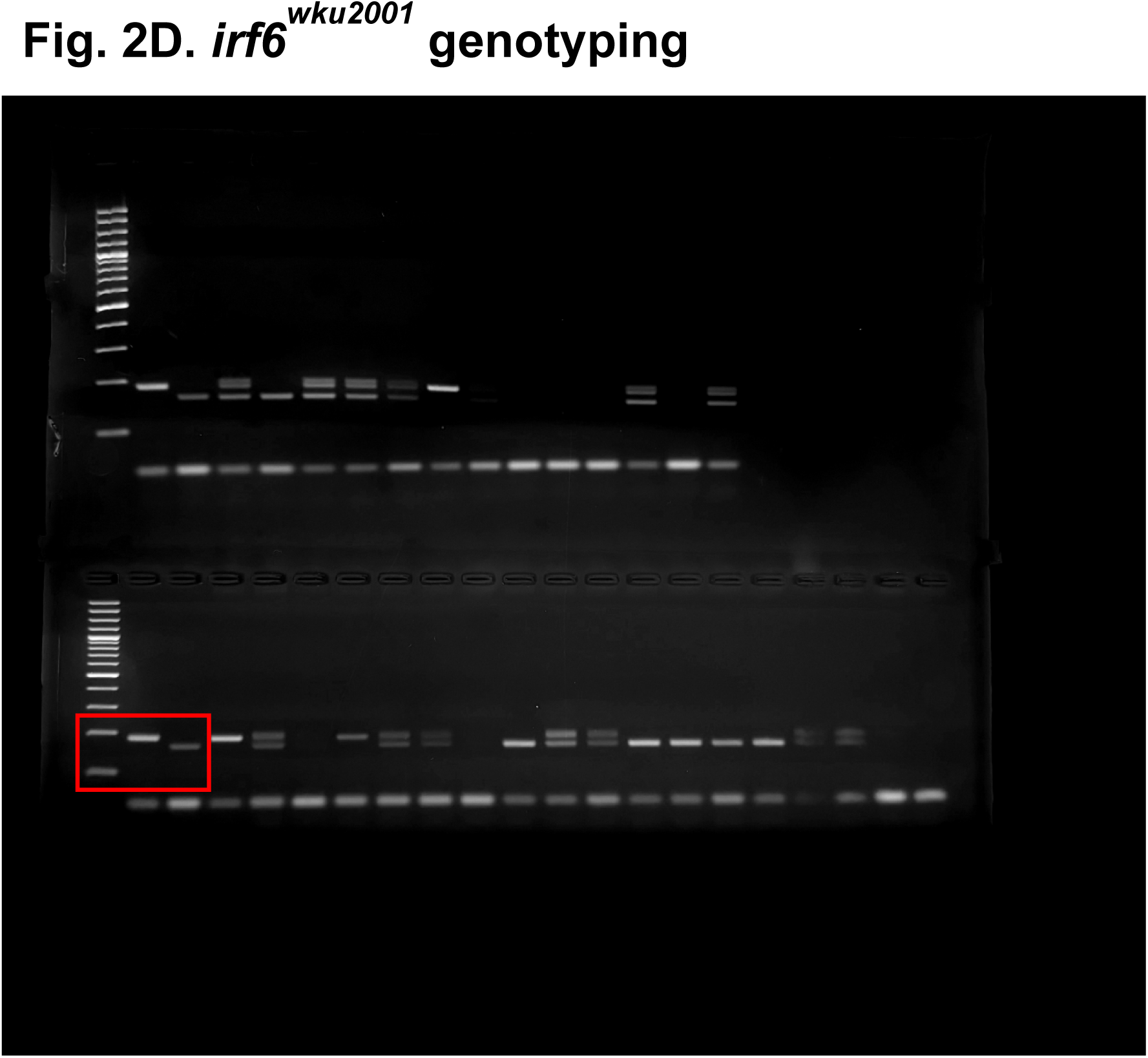

